# Early Cretaceous Origin and Evolutionary History of Palms (Arecaceae) inferred from 1,033 Nuclear Genes and a New Synthesis of Fossil Evidence

**DOI:** 10.1101/2024.06.23.600266

**Authors:** Sidonie Bellot, Fabien L. Condamine, Kelly K.S. Matsunaga, Robert J. Morley, Ángela Cano, Thomas L.P. Couvreur, Robyn Cowan, Wolf L. Eiserhardt, Benedikt G. Kuhnhäuser, Olivier Maurin, Michelle Siros, Felix Forest, Ilia J. Leitch, William J. Baker

## Abstract

Tropical rainforests are home to almost half of all known plant diversity, yet a shortfall in tropical plant phylogenetic hypotheses is hindering our understanding of how rainforests have formed and adapted to past global changes. Phylogenetic and historical biogeographic evidence from key rainforest lineages, including palms (Arecaceae), are required to illuminate the history of these ecosystems. However, our current understanding of the palm tree of life is based on uneven sampling of data from the plastid and nuclear genome, and numerous palm genera and palm fossils have been described or revised over the past decade, casting doubt on many palm relationships, ages and ancestral ranges inferred in previous studies. Here, we infer the phylogenetic relationships of all palm genera based on data from 1,033 nuclear genes generated using target sequence capture. Our palm tree of life is highly resolved and supported. Remaining areas of ambiguity reflect the complex dynamics of palm evolution, including many rapid diversification events in subfamily Arecoideae and putative cases of ancient reticulation throughout the family. We review the phylogenetic placement and age of hundreds of palm fossils and use a vetted selection to estimate divergence times and ancestral ranges for all palm genera. We show that the family is older than previously thought, likely first diversifying in the Early Cretaceous in Laurasia, and that three quarters of the genera had originated by the Oligocene. The early spread of palms across the world involved at least 40 long-distance dispersals across oceanic gaps. Dispersals away from northern latitudes vastly outnumber dispersals into these latitudes, consistent with the ancient affinity of palms for megathermal climates, and suggesting that global cooling during the Oligocene was a major constraint on early palm distribution. Our dated phylogenomic tree and curated fossil dataset provide a new foundation for evolutionary studies on palms, opening the door to deeper research on the rainforest biome in which they thrive.

## Introduction

The biological megadiversity of the tropical rainforests is increasingly threatened, yet the mechanisms underpinning this diversity, its evolution and adaptations to environmental changes remain poorly understood. Phylogenetic trees are the cornerstone of studies addressing rainforest diversity through comparative trait studies, divergence time estimates, and biogeographical inferences. However, there is a shortfall in phylogenetic information from the tropics, notably in plants (Rudbeck et al., 2022). This knowledge gap demands urgent attention, especially for rainforests, which contain *ca.* 45% of global plant species (Eiserhardt et al., 2017), many of which are at risk of extinction (Pelletier et al., 2018).

The palm family (Arecaceae) is a well-studied, keystone component of tropical rainforests with a wealth of biological data supporting its establishment as a model group for rainforest research (Couvreur et al., 2011; Couvreur & Baker, 2013; Dransfield et al., 2008; Kissling, Eiserhardt, et al., 2012). These include a complete genus-level phylogenetic tree (Baker et al., 2009), comprehensive taxonomic and geographic data (Govaerts et al., 2021; Govaerts & Dransfield, 2005), and a well-documented fossil record (Harley, 2006). These resources have enabled a series of studies providing insights into palm biogeography and diversification, and the origins and evolution of rainforests (Baker & Couvreur, 2013a, 2013b; Couvreur et al., 2011, 2015; Kissling, Baker, et al., 2012; Kissling, Eiserhardt, et al., 2012). However, over the last decade, DNA sequencing techniques have changed markedly, new fossils have emerged and biogeographically relevant taxa have been discovered (see below), casting new light on earlier research. In view of the significance of these developments, a new synthesis of palm evolution and biogeography is now needed.

Family-wide phylogenetic studies of palms date back to the mid-1990s, commencing with analyses of plastid restriction fragment length polymorphisms (Uhl et al., 1995), followed by a series of Sanger sequencing studies, reporting an increasingly comprehensive sampling for the plastid regions *trnL–trnF*, *rps16* intron, *rbcL* and *matK* (Asmussen et al., 2000, 2006; Asmussen & Chase, 2001; Baker et al., 1999). These studies were integrated with other plastid and nuclear gene datasets (Baker et al., 2000; B. F. Gunn, 2004; Hahn, 2002a, 2002b; Lewis & Doyle, 2001, 2002; Loo et al., 2006; Norup et al., 2006; Roncal et al., 2005; Savolainen et al., 2006; M. M. Thomas et al., 2006) and a novel morphological dataset to produce the first phylogenetic tree to include all palm genera recognized at the time (Baker et al., 2009). This study not only provided the foundations on which a new phylogenetic classification of palms was built (Dransfield et al., 2008) but also led to much downstream evolutionary research (e.g.: Antonelli et al., 2017; Cássia-Silva et al., 2019; Faurby et al., 2016; Onstein et al., 2017; Torres Jiménez et al., 2023). However, the classification of palms has subsequently evolved, with some genera split, others merged and new ones discovered as new-to-science (Baker & Dransfield, 2016; Eiserhardt et al., 2022; Hodel et al., 2021; Pérez-Calle, Bellot et al., 2024; Sâm et al., 2023), rendering the existing synthesis incomplete. Moreover, phylogenomic methods have entirely superseded previous approaches applied to palms, through plastome sequencing (Barrett et al., 2016; Chen et al., 2022; Comer et al., 2015; Yao et al., 2023) and targeted sequencing of nuclear genes (Cano et al., 2022; Comer et al., 2016; de La Harpe et al., 2019; Eiserhardt et al., 2022; Escobar et al., 2022; Helmstetter et al., 2020; Heyduk et al., 2016; Kuhnhäuser et al., 2021; Loiseau et al., 2019; Pérez-Calle, Bellot et al., 2024; Sanín et al., 2022; Torres Jiménez et al., 2021). Notable among these is the near-complete, genus-level, plastome-based study of Yao et al. (2023). A nuclear phylogenomic, palm tree of life is now required to integrate the complex and rich phylogenetic signal inherent in nuclear genes. The availability of lineage-specific (de La Harpe et al., 2019; Heyduk et al., 2016; Loiseau et al., 2019) and universal (Johnson et al., 2019) target sequence capture probes has brought this goal well within reach.

The spatio-temporal framework for palm diversification (Baker & Couvreur, 2013b, 2013a; Couvreur et al., 2011) established through molecular dating of the genus-level tree of Baker et al. (2009) challenged prevailing understanding of both palm biogeography and rainforest biome evolution. Four fossils were used to calibrate the tree, selected on the basis of high confidence in their taxonomic affinities and reported ages (Couvreur et al., 2011). However, the fossil record of palms has received a surge of attention in recent years (e.g.: Bogotá-Ángel et al., 2021; Collinson et al., 2012; Greenwood et al., 2022; Jaramillo et al., 2014; Kumar et al., 2022; Lim et al., 2022; Manchester et al., 2016; Martínez et al., 2016; Matsunaga & Smith, 2021; Parmar et al., 2023; Shukla et al., 2012; Singh et al., 2016). It is unrivalled among monocotyledonous plant families (Iles et al., 2015), with numerous records from the Late Cretaceous (Dransfield et al., 2008; Harley, 2006), and some that might extend into the Early Cretaceous (Martínez et al., 2016). The palm fossil record now requires re-assessment for its application in molecular dating to account for new research and discoveries that may lead to substantial changes in inferred dates, and to capitalise on biogeographic methods that can incorporate data from the fossil record explicitly.

Here, we present a new genus-level tree of life for the palm family. Using two distinct target capture probe sets, we sequenced up to 1,033 nuclear genes from all currently accepted palm genera and inferred their phylogenetic relationships, highlighting potential cases of rapid divergence and reticulation. We then reviewed the palm fossil record to establish a vetted set of fossils for molecular dating and biogeographical inferences. We used a selection of these fossils to estimate divergence times and ancestral ranges. We then exploited this spatio-temporal framework to build a comprehensive picture of the early diversification of palms and their rise to prominence across the tropics.

## Materials & Methods

### Taxon sampling, DNA Extraction and Sequencing

Leaf tissue from silica-dried or herbarium specimens was sampled for at least one species from each of the 184 genera currently recognized in the family (Baker & Dransfield, 2016; Eiserhardt et al., 2022; Hodel et al., 2021; Pérez-Calle, Bellot et al., 2024; Sâm et al., 2023) and from the outgroup genus *Dasypogon* (Dasypogonaceae; Barrett et al., 2016; Li et al., 2021). See Supplementary Methods and Table S1 for details. DNA was extracted using a modified cetrimonium bromide (CTAB) method (Doyle & Doyle, 1987) as described in Brewer et al. (2019). Dual indexed DNA libraries were prepared using the DNA NEBNext® Ultra™ II Library Prep Kit from New England BioLabs (Ipswich, MA, USA) and NEBNext® Multiplex Oligos for Illumina® (Dual Index Primers Sets 1 and 2), following the manufacturer’s protocols, including a fragmentation step when necessary (Supplementary Methods). Pools of up to 48 DNA libraries were enriched in selected genes using two Daicel Arbor Biosciences myBaits® probe kits: Angiosperms353 (Johnson et al., 2019) and PhyloPalm (Loiseau et al., 2019), targeting 353 and 970 nuclear genes shown to be phylogenetically informative across angiosperms and palms, respectively. Hybridisation and Illumina paired-end DNA sequencing were conducted separately for each probe kit (Supplementary Methods).

### Read Processing, Sequence Assembly and Alignment

Sequence data quality was assessed using FASTQC v. 0.11.9 (Andrews, 2010), and Trimmomatic v. 0.39 (Bolger et al., 2014) was used to remove adapters and bases with low quality (Supplementary Methods). Clean reads were assembled using HybPiper v. 1.3 (Johnson et al., 2016) to recover and assemble the regions targeted by both probe kits (i.e. 1,255 non-overlapping regions, see Supplementary Methods). To mitigate the risk of analysing paralogs, we expanded an approach first developed for subfamily Calamoideae (Kuhnhäuser, 2021) by classifying the target regions into orthologous vs. paralogous regions based on their copy numbers in the annotated genomes of palm species representing the three main palm subfamilies Arecoideae, Calamoideae and Coryphoideae (Supplementary Methods). This resulted in the classification of 951 regions as paralogous and 304 as orthologous. For each region, sequences shorter than 250 bp and/or than 25% of the median sequence length for the region were discarded to reduce the risk of alignment errors (Smirnov & Warnow, 2021). Sequences were then aligned with MAFFT v. 7 (Katoh & Standley, 2013), and alignments were cleaned using OptrimAl (https://github.com/keblat/bioinfo-utils/blob/master/docs/advice/scripts/optrimAl.txt), CIAlign v. 1.1.0 (Tumescheit et al., 2022), and TAPER v. 1.0 (Zhang et al., 2021) (Supplementary Methods), yielding 1,118 clean alignments. Because missing taxa can lead to errors in species tree inference (Nute et al., 2018), gene trees were computed only from the 1033 alignments comprising at least 75 samples (40%), 231 of which were derived from regions classified as orthologous.

### Phylogenetic Inferences and Branch Support Evaluation

The model of nucleotide substitution best fitting each alignment was estimated using the ModelFinder (Kalyaanamoorthy et al., 2017) method implemented in IQ-TREE v. 1.6.12 (Minh et al., 2020). A maximum likelihood (ML) tree (hereafter referred to as “gene tree”) was then estimated from each alignment using RAxML-ng v. 1.1 (Kozlov et al., 2019), setting the model of substitution to the one identified by the IQ-TREE analysis and performing 60 searches based on 30 parsimony-based randomised trees and 30 random trees. Branch support was estimated based on 500 bootstrap replicates. After collapsing branches with less than 10% bootstrap support using newick-utils v. 1.6 (Junier & Zdobnov, 2010), ASTRAL-III v. 5.7.5 (Zhang et al., 2018) was used to generate a species tree from the gene trees. A second species tree was generated using orthologous regions only.

Branch support was assessed using local posterior probability (LPP) and quartet scores provided by ASTRAL. Branches were classified as resolved with strong support if their LPP was ≥ 0.90 and as resolved with weak support otherwise, following Sayyari & Mirarab (2016). For each internal branch of the species tree, we tested whether or not the two alternative topologies appeared in statistically different frequencies among the gene trees, which would suggest that the observed gene tree discordance may not only be due to incomplete lineage sorting (ILS) or gene tree error (GTE), but also to other phenomena such as introgression. Topology frequencies were compared using a binomial test with and without Bonferroni correction for multiple testing across the phylogenetic tree (p-value thresholds of 0.00027 and 0.05, respectively).

### Fossil Literature Review

We reviewed the palm fossil literature, building on existing pollen and macrofossil reviews (Bogotá-Ángel et al., 2021; Dransfield et al., 2008; Harley, 2006; Harley & Baker, 2001; Iles et al., 2015; Lim et al., 2022). We searched specifically in the scientific literature for fossils described or revisited after 2008, which corresponds to the date of the most recent comprehensive palm macrofossil review (Dransfield et al., 2008). We prioritised the compilation of fossils that could be assigned to a palm subfamily and to a geological period, epoch or stage as these were most informative for calibrating phylogenetic trees. Broadly following the recommendations of Parham et al. (2012), phylogenetic affinity and age reliability was assessed for each fossil (or group of fossils from the same time, place and lineage) based on the literature and our own expertise, adding comments when our conclusions differed from the original assignments.

The complete list of fossils and with referenced taxonomic, phylogenetic, geographical, and age information, can be found in Table S2. Our list contained 194 records for macrofossils, including nine assigned to Arecoideae, 51 to Coryphoideae, 14 to Nypoideae, two to Calamoideae, 80 not assigned to a subfamily, and 38 previously assigned to palms that were here revised as “possibly Arecaceae” in the absence of substantial evidence that they could not be anything else than a palm. For pollen, the list included 104 records, including 14 assigned to Arecoideae, six to Coryphoideae, eight to Nypoideae, one to Ceroxyloideae, 66 to Calamoideae, eight not assigned to a subfamily, and one considered “possibly Arecaceae”. We grouped individual fossil reports in a single record if they belonged to the same taxon, geological horizon and region (i.e. North Africa, Central Africa, South Africa, Europe, Africa, India, Indian Ocean, Madagascar, Asia, Oceania; see definitions in Supplementary Methods), while providing details about finer geographical distribution for each record (Table S2).

### Molecular Dating Analyses

Divergence times were estimated using an optimised relaxed clock (ORC) model (Douglas et al., 2021) implemented in BEAST 2.6 (Bouckaert et al., 2019), based on 22 orthologous regions selected using SortADate (Smith et al., 2018) and concatenated using AMAS (Borowiec, 2016). Detailed parameter settings are provided in Supplementary Methods. Analyses were performed with and without constraining the tree topology based on the ASTRAL species tree obtained from summarising all the genes. Number of generations and effective sampling sizes were assessed using Tracer v. 1.7.2 (Rambaut et al., 2018) and summarised in Table S3. Maximum clade credibility trees with median node ages were generated from the post-burn-in posterior tree distribution of each analysis using TreeAnnotator, which is part of BEAST.

We calibrated the minimum age of six internal nodes using six fossils distributed across the palm family (details in Supplementary Methods; geological periods follow Cohen et al. (2013): 1) *Mauritiidites* spp. (stem of Mauritiinae; pollen; Maastrichtian to Campanian of Nigeria, Sudan, Egypt, Cameroon, Gabon and Angola; Edet & Nyong, 1993; Eisawi & Schrank, 2008; Rull, 1998; Salami, 1990; Salard-Cheboldaeff, 1990; Schrank, 1994); 2) *Dicolpopollis* spp. (stem of Metroxylinae + Plectocomiinae + Calaminae + Pigafettinae; pollen; Cenomanian of Australia; Burger, 1990; Macphail & Jordan, 2015; Totterdell & Mitchell, 2009); 3) *Sabal bigbendense* Manch., Wheeler, & Lehman (stem of *Sabal*; seed; Campanian of Texas; A. Cano, 2018; Manchester et al., 2010); 4) *Hyphaeneocarpon indicum* Bande, Prakash, & Ambwani emend.

Matsunaga, S.Y.Sm., Manch., Srivastava, & Kapgate (crown of Hyphaeninae; fruit; Paleocene of India; Bande et al., 1982; Matsunaga et al., 2019); 5) *Palmocarpon drypetoides* (Mehrotra, Prakash & Bande) Manch., Bonde, Nipunage, Srivastava, Mehrotra & S.Y.Sm. (crown of Attaleinae; fruit; Paleocene of India; Manchester et al., 2016) and 6) *Friedemannia messelensis* Collinson, Manch. & Wilde (crown of Areceae; fruit; Eocene of Germany; Collinson et al., 2012; Matsunaga & Smith, 2021). The placement of these last three fossils was supported by previous morphology-based phylogenetic inference (Matsunaga & Smith, 2021). However, given the relatively young age and limited number of characters supporting the placement of *F. messelensis* at the crown node of tribe Areceae (Matsunaga & Smith, 2021), we performed analyses with and without this fossil. The prior age distributions and maximum ages of the six calibrated nodes were defined to account for the well-known issue that fossils may not be the oldest representatives of a lineage (Magallón, 2004).

Following strategies adopted in recent angiosperm-wide studies (Ramírez-Barahona et al., 2020; Zuntini, Carruthers et al., 2024), the priors followed a log-normal distribution with an offset corresponding to the fossil age, and the mu and sigma parameters set so that the 97.5% quantile of the node age would correspond to the 97.5% quantile of the root age prior distribution (see below).

Two approaches were used to calibrate the root age (i.e. the stem node of palms). The first (referred to as the “younger ages” analysis), followed the rationale of Couvreur et al. (2011) that palms are unlikely to be older than the oldest known monocot fossils, which are *ca.* 113 Ma old (Coiffard et al., 2019; Iles et al., 2015), and used a normal root age prior distribution with a mean of 113 Myr and a standard deviation of 2. The second approach (referred to as the “older ages” analysis) accommodated the possibility that palms, or indeed monocots, may be much older than their oldest reliably dated fossils, and used instead a normal root age prior distribution with a mean of 149.1 Myr and a standard deviation of 28. This was chosen so that the lower bound corresponded to the age of the oldest palm fossil assigned to a subfamily (calibration 2 above) and so that the upper bound would be in line with the maximum age obtained in previous angiosperm-wide studies for this node (Ramírez-Barahona et al., 2020; Zuntini, Carruthers et al., 2024). The two sets (younger and older ages) of four analyses (with and without *F. messelensis* and with and without constraining the tree topology) are summarised in Table S3. For a given strategy, analyses with or without constraining the tree topology provided very similar ages, as did analyses using or not *Friedemannia messelensis* to calibrate the age of crown Areceae (Fig. S1), so downstream analyses and results only rely on the “younger” and “older” dated trees obtained by excluding *F. messelensis* from the calibrations and by constraining the tree topology.

### Historical Biogeography Inferences

Palm ancestral ranges were estimated using the Dispersal-Extinction-Cladogenesis (DEC) model (Ree & Sanmartín, 2018), implemented in DECX (Beeravolu & Condamine, 2016; available at: https://github.com/champost/DECX; Supplementary Methods). Connectivity between areas was constrained based on how distances between areas changed through geological periods (Supplementary Methods). Extant and fossil taxa distribution ranges were coded by dividing the world into ten areas: North America, Central America, South America, Europe, Asia, Africa, Oceania, India, Indian Ocean, and Madagascar. Another coding reproducing the approach of (Baker & Couvreur, 2013a) was also used, comprising seven areas: North America (including Central America) South America, Africa, India, Eurasia, Oceania and Indian Ocean (including Madagascar). The delimitation between Asia and Oceania followed the Lydekker line in the 10-area scheme, and the Wallace line in the 7-area scheme (Supplementary Methods). The distribution range of the species included in the tree was coded in two ways, the first based on the distribution of the whole genus to which they belonged, the second based on the distribution of the sampled species (POWO, 2024; Table S1). The distributions were the same in both codings for 77%-78.6% of the species, depending on the area classification (Table S1). Ancestral biogeographic ranges spanning more than four areas were only allowed if they were subsets of the tip ranges.

Biogeographic analyses were performed on the “younger” and “older” dated trees. For each tree, they were done with and without using information about the location of four (older tree) to eight (younger tree) fossils to inform ancestral ranges (Supplementary Methods). This resulted in 16 ancestral range inferences (younger vs older tree * with vs. without fossil information * first vs. second area coding * species vs. genus-level range coding), which are further described in Supplementary Methods. Dispersals between the biogeographic areas were visually inferred based on the ancestral range estimates.

## Results

### Genus-Level Phylogenetic Relationships within Palms

Phylogenetic analyses based on 1,033 (all-genes tree; Fig. 1) or 231 nuclear genes (orthologs-only tree; Fig. S2) supported the division of Arecaceae into the five currently accepted subfamilies with Calamoideae, the monotypic Nypoideae and Coryphoideae together forming a grade around sister subfamilies Ceroxyloideae and Arecoideae. The monophyly of all non-monotypic subfamilies, tribes and subtribes was recovered with strong support (LPP ≥ 0.9) or, in the case of Archontophoenicinae and Rhopalostylidinae, with weak support (LPP < 0.9). There were two exceptions in subtribe Areceae: *Calyptrocalyx* was excluded from Laccospadicinae and resolved instead as sister to Archontophoenicinae with strong support, and Rhopalostylidinae were nested in Basseliniinae with weak support (Figs. 1, S2). The all-genes and orthologs-only trees were both strongly supported and highly congruent: 168 (91%) branches were identically resolved in both trees, including 139 strongly supported in both trees, while only 17 (9%) branches were resolved differently, none of them with strong support in both trees (Fig. S2).

**Figure 1.**
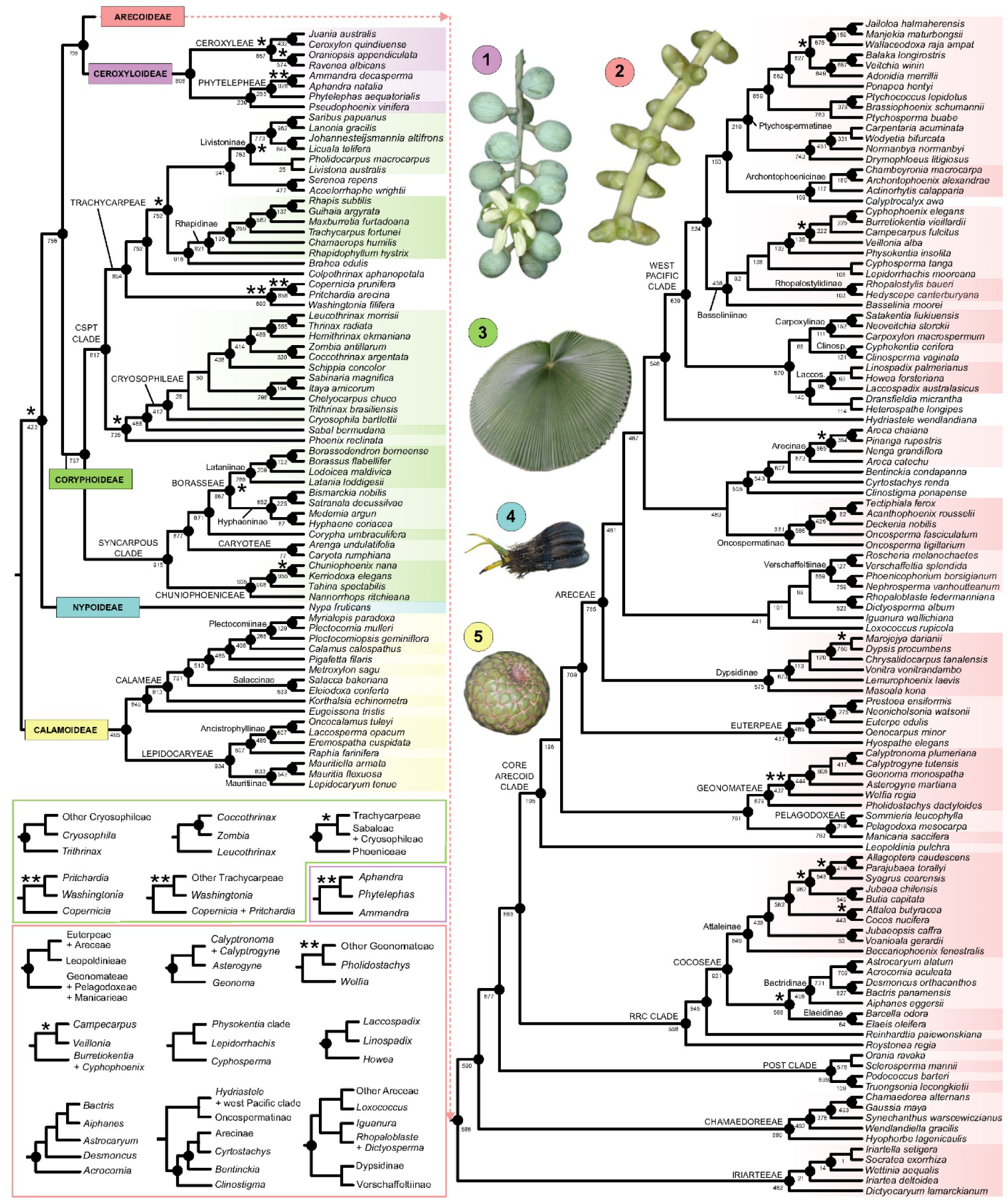
Phylogenetic relationships among palm genera based on 1,033 nuclear genes (all-genes tree). Circles at nodes indicate strong support, corresponding to local posterior probabilities ≥ 0.9. Stars indicate branches significantly associated with ILS/GTE-inconsistent discordance without (*) or with (**) Bonferroni correction (see Methods). Numbers below branches indicate the number of gene trees in which that branch was recovered. **Inset: Alternative topologies for branches different in the orthologs-only tree and for selected branches associated with ILS/GTE-inconsistent discordance.** For branches differing between the all-genes and orthologs-only trees, the orthologs-only topology is displayed. For branches that were the same in both trees but were associated with ILS/GTE-inconsistent discordance after Bonferroni correction was applied (**), the most frequent alternative topology found among the gene trees is displayed. Pictures show key characteristics of each subfamily with 1: solitary flowers of *Pseudophoenix vinifera* (Ceroxyloideae); 2: triads, each comprising two male and one female flower buds, in *Manjekia maturbongsii* (Arecoideae); 3: fan leaf of *Licuala cordata* (Coryphoideae); 4: seed of *Nypa fruticans* germinating viviparously (Nypoideae); 5: fruit of *Metroxylon sagu* covered with scales (Calamoideae). All pictures taken by W. J. Baker.

The average number of genes underpinning strongly supported branches was significantly higher than for weakly supported branches (t-test p-values < 0.001), yet some strongly supported branches were underpinned by few genes (Fig. S3). The trees are described in detail in Supplementary Results & Discussion. The relationships of several genera of tribe Trachycarpeae that are unplaced at the subtribe level in the current classification due to past uncertainty (Baker & Dransfield, 2016) were resolved with strong support: *Brahea* as sister to Rhapidinae; *Serenoa* and *Acoelorrhaphe* forming a clade sister to Livistoninae, and *Colpothrinax* as sister to all Trachycarpeae except *Pritchardia*, *Copernicia* and *Washingtonia*. Similarly in tribe Areceae, strongly supported relationships were recovered for genera previously unplaced to subtribe: *Rhopaloblaste* and *Dictyosperma* were strongly supported as sister genera, *Clinostigma*, *Cyrtostachys* and *Bentinckia* formed a strongly supported grade in which Arecinae was nested, and *Hydriastele* was sister to the “west Pacific clade” (described in Baker et al., 2011), in which *Dransfieldia* and *Heterospathe* formed a clade (Fig. 1). However, the placement of the other unplaced genera of Areceae (*Iguanura*, *Bentinckia* and *Loxococcus*) were different and weakly supported in both trees (Figs. 1, S2).

Gene tree discordance inconsistent with ILS or GTE (see Methods) was found to occur at four branches (two involving the placements of *Pritchardia*, *Copernicia* and *Washingtonia,* one involving *Welfia,* and one in Phytelepheae), all strongly supported and identically resolved in both trees (Fig. 1). This number increased to 22 branches when not applying a Bonferroni correction (Figs. 1, S2; Supplementary Results & Discussion). The median number of genes informing the resolution of branches affected by ILS/GTE-inconsistent discordance was higher than for branches not affected by it (Fig. S3).

### A Curated Fossil Set for Palm Molecular Dating and Biogeography

The distribution of palm fossil records across time and space for all Arecaceae and for the three largest subfamilies (Arecoideae, Calamoideae and Coryphoideae) is visualised in Figure 2. Arecoideae showed similar numbers of records for macro- and pollen fossils, while most entries for Coryphoideae were macrofossils and most entries for Calamoideae were pollen records (Fig. 2).

**Figure 2.**
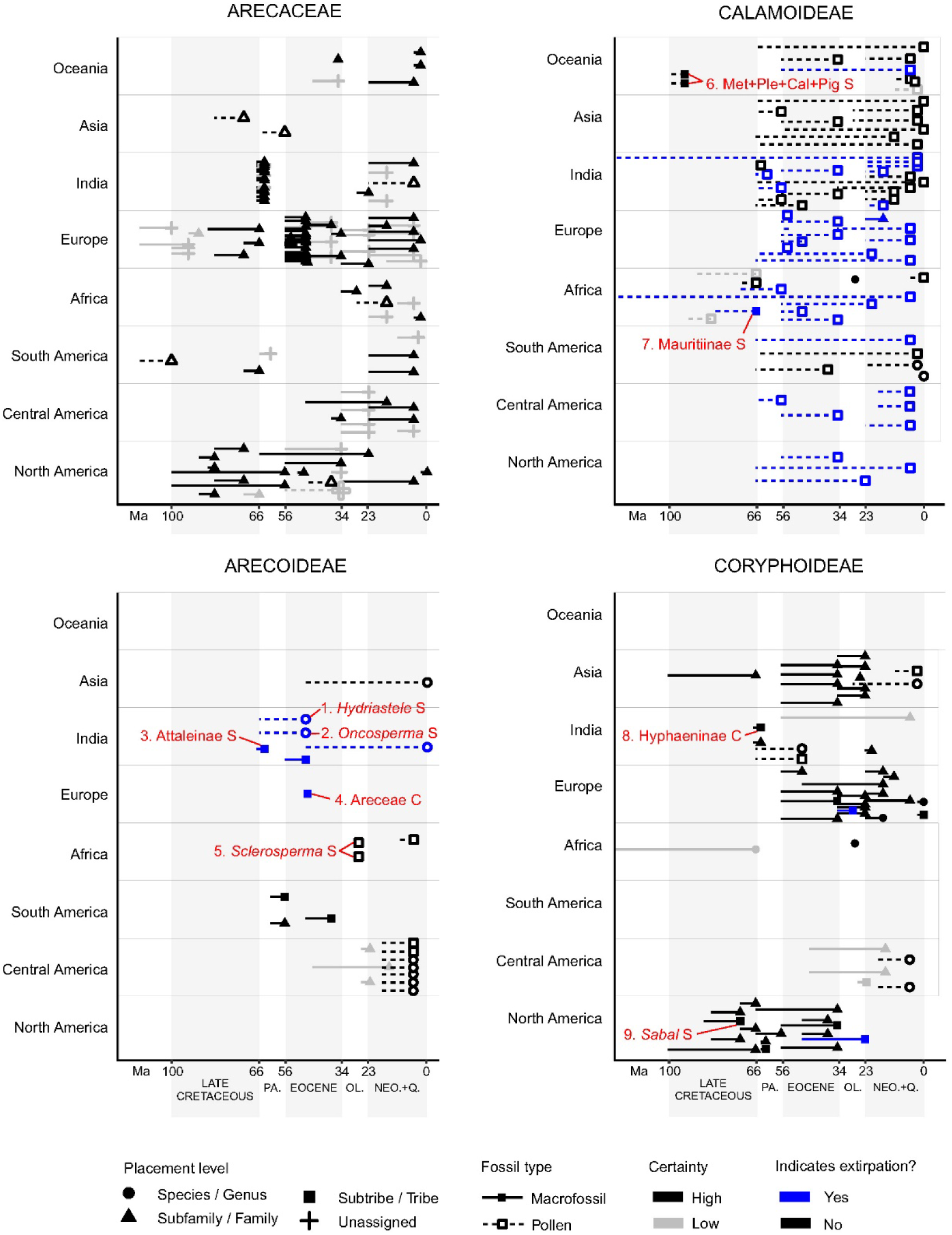
Global records of fossils assigned to Arecaceae, Arecoideae, Calamoideae and Coryphoideae from the Early Cretaceous to the Quaternary. All fossil records are described in Table S2. The Arecaceae panel only comprises fossils that could not be placed below the family level. Grey bars and symbols are used when the fossil age and/or its phylogenetic placement was considered uncertain during our review. Details regarding the uncertainty are provided in Table S2. When the fossil records are known from a given time period without further precision, the bar spans the entire period. Extirpation is here understood as a mismatch between the area where the fossil was recorded and the current distribution range of the lineage to which it was assigned; such cases are of high interest to inform ancestral range inferences. Fossils that can be used to calibrate minimum clade ages are annotated with the name of the clade (S: stem; C: crown; Met+Ple+Cal+Pig: Metroxylinae + Plectocomiinae + Calaminae + Pigafettinae), and the fossil names are as follows: 1: *Retiminocolpites perforatus*; 2: *Clavapalmaedites clavatus*; 3: *Palmocarpon drypeteoides*; 4: *Friedemannia messelensis*; 5: *Sclerosperma* spp.; 6: *Dicolpopollis* spp.; 7: *Mauritiidites* spp.; 8: *Hyphaeneocarpon indicum*; 9: *Sabal bigbendense* and *Sabal bracknellense* (details in Table S2). P.: Paleocene; EOC.: Eocene, OL.: Oligocene; NEO. + Q.: Neogene + Quaternary.

Numerous Calamoideae (mainly pollen) and Nypoideae fossils were found across all regions (Fig. 2, Table S2). Arecoideae fossils were mainly found in India, Africa, and Central and South America. Coryphoideae fossils were mainly found in North America, Europe, India and Asia. Most fossils that could not be assigned to a subfamily or that could not be confirmed as palms were from Europe, the Deccan Intertrappean Beds (India) and North America (Fig. 2).

Some fossils were identified as being of strong interest for biogeographic analysis because they were discovered in areas in which the clade they represent is not currently found (Table S2). This included 1) European fossils from the Eocene, Oligocene, and Miocene assigned to Arecoideae (Areceae; Chandler, 1957; Collinson et al., 2012; Matsunaga & Smith, 2021), Nypoideae (Gee, 1990; Haseldonckx, 1972), and Calamoideae (including Calameae, Calaminae, Salaccinae, Mauritiinae and Eugeissoneae; e.g.: Akkiraz et al., 2006, 2008; Bogotá-Ángel et al., 2021; Czeczott & Juchniewicz, 1975; Filatoff & Hughes, 1996; Hofmann et al., 2015; Jablonszky, 1914; Lim et al., 2022; Table S2), 2) a Cryosophileae (Coryphoideae) fossil stem from the Oligocene of Europe (R. Thomas & De Franceschi, 2012), 3) North American fossils assigned to Calamoideae (Calameae, Calaminae, and Eugeissoneae; Lim et al., 2022; Tschudy, 1973) and to *Phoenix* (Coryphoideae; E. W. Berry, 1914, 1924), 4) Eugeissoneae fossils from Africa, Oceania, Central and South America (Lim et al., 2022), 5) *Korthalsia*, *Eugeissona*, Mauritiinae (Calamoideae; Lim et al., 2022; Parmar et al., 2023; Phadtare & Kulkarni, 1984; Ramanujam et al., 1991, 1992; Sarma et al., 1984), *Cocoseae*, *Hydriastele*, and *Oncosperma* (Arecoideae) fossils from India (Guleria et al., 1996; Manchester et al., 2016; Matsunaga et al., 2019; Parmar et al., 2023; Shukla et al., 2012; Singh et al., 2016), 6) Mauritiinae fossils from Africa and Central America (Bogotá-Ángel et al., 2021; Edet & Nyong, 1993; Eisawi & Schrank, 2008; Jaramillo et al., 2014; Okeke & Umeji, 2016; Rull, 1998; Salard-Cheboldaeff, 1990; Schrank, 1994; Srivastava & Binda, 1991), and 7) Nypoideae fossils found in Africa (e.g.: Bonnet, 1904; Gregor & Hagn, 1982) and the Americas (e.g.: E. Berry, 1914, 1916; Gomez-Navarro et al., 2009; Wing et al., 2009) until the Miocene (Lim et al., 2022; Table S2).

The minimum age of most palm fossils fell between the Miocene and the Paleocene but some dated as Cretaceous. The earliest records for key clades were identified as potentially useful for calibrating the minimum age of nodes when dating palm phylogenetic trees (Fig. 2; Table S2). The earliest confirmed record for the whole palm family was pollen dating at least from the Albian (100.5–113 Ma) of Patagonia (Argentina) with unclear phylogenetic placement, perhaps representing a lineage ancestral to Calamoideae or to all palms (Martínez et al., 2016; Table S2).

This was followed by Calamoideae pollen records from the Cenomanian of Australia, some of which were assigned to a clade nested inside tribe Calameae (Burger, 1990; Macphail & Jordan, 2015; Totterdell & Mitchell, 2009; Fig. 2; Table S2) and others to tribe Eugeissoneae (Table S2). Other early records included macrofossils from the Turonian of Europe (Crié, 1892; Kvaček & Herman, 2004), the Santonian and/or Coniacian of North America (E. Berry, 1916; Greenwood et al., 2022) and the Campanian of Asia (Harley, 2006; Takahashi, 1964) that could not be assigned below family level.

The earliest Coryphoideae were identified as *Sabal* and *Sabalites* fossils from the Campanian of North America (E. Berry, 1914b; A. Cano, 2018; Greenwood et al., 2022; Manchester et al., 2010; Fig. 2), while the earliest Arecoideae were recorded from the Deccan Intertrappean Beds of India (64-67 Ma; Manchester et al., 2016; Matsunaga & Smith, 2021; Fig. 2). In Nypoideae, pollen was recorded from the Late Cretaceous of Asia, India, Oceania, and South America, and more specifically from the Campanian of Africa (Eisawi & Schrank, 2008; Gee, 1990; Lim et al., 2022; Table S2). Beyond these earliest records across subfamilies, younger records were identified as potentially suitable for age calibration of younger nodes (Fig. 2). These included an Areceae fruit from the Eocene (Collinson et al., 2012; Matsunaga & Smith, 2021; Fig. 2), pollens from the Lower Eocene to Paleocene assigned to *Hydriastele* and *Oncosperma* (Parmar et al., 2023; Fig. 2), *Sclerosperma* pollen from the Oligocene (Grímsson et al., 2019; Fig. 2), Mauritiinae pollen from the Santonian and possibly Coniacian (Atta-Peters & Salami, 2006; Bogotá-Ángel et al., 2021; Edet & Nyong, 1993; Fo & Fa, 2018; Fig. 2; Table S2), and a Hyphaeninae fruit from the Deccan Intertrappean beds (Bande et al., 1982; Matsunaga & Smith, 2021; Fig. 2).

### Palm Divergence Times and Ancestral Distribution Ranges

The median ages obtained with the “younger ages” strategy (see Methods) were 21 Myr younger on average compared to those obtained with the “older ages” strategy (Fig. 3, Fig. S4, Table S4). Irrespective of the molecular dating strategy, palms were inferred to have originated by the end of the Early Cretaceous, and most genera (74-92%) were inferred to have originated by the end of the Oligocene (23 Ma), including 45-76% by the end of the Eocene (33.9 Ma), while only 8-26% (mostly in the west Pacific clade of tribe Areceae) originated in the Miocene (23–5 Ma; Fig. S4). Genus-level, 10-area coding, ancestral range inferences based on the younger ages indicate that the most recent common ancestor (MRCA) of palms occurred 119 Ma (95% highest posterior density interval: 116–123 Ma; Figs. 3, S4; Table S4) in Laurasia and Central America, and possibly also in South America according to alternative states with similar probabilities (Figs. 3, S5; Table S5). Palm subfamilies appear to have all diverged from each other by 101 Ma (95–107 Ma; Figs. 3, S4; Table S4). The MRCA of Calamoideae was estimated to be 107 (103–112) Myr old and its range most likely comprised North America and Asia, and possibly already Europe and Central America, where the subfamily was present by the end of the Cretaceous (Figs. 3, S4-S7). Nypoideae diverged from the three other subfamilies 112 (106–116) Ma, and their MRCA most likely occurred in Asia, and possibly also in America (Figs. 3, S4-S7). The MRCA of Coryphoideae most likely occurred in Asia and North America 100 (93–105) Ma (Figs. 3, S4-S7), and that of Ceroxyloideae appeared to have occurred in Central, and possibly North, America 78 (61–94) Ma (Figs. 3, S4-S7). Finally, the MRCA of Arecoideae was inferred to have occurred in Asia and Central America, and possibly also in North and South America 101 (95–107) Ma (Figs. 3, S4-S7).

**Figure 3.**
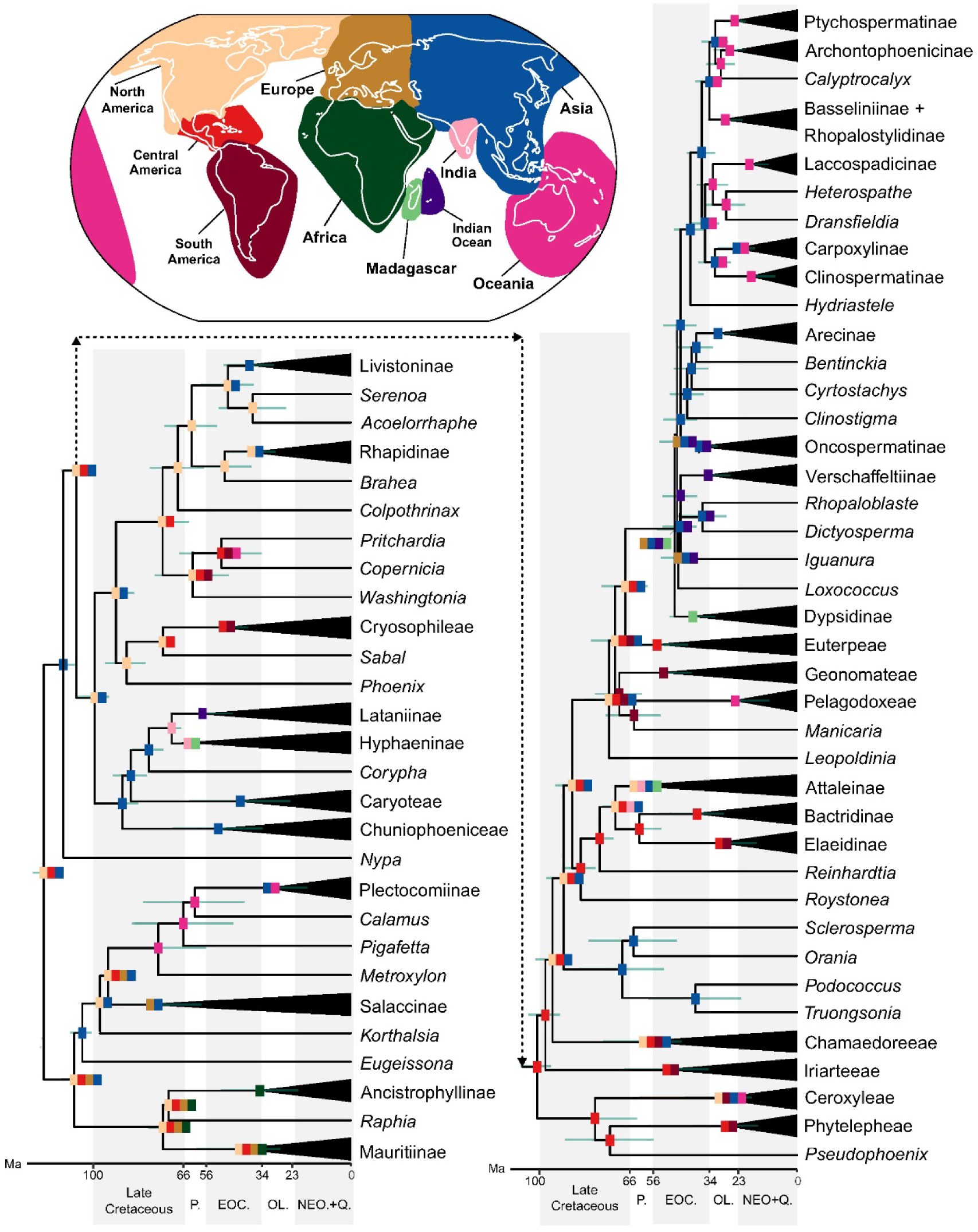
Palm ages and ancestral distribution ranges. The tree shows palm phylogenetic relationships with non-monotypic subtribes collapsed and palm median ages obtained from the “younger ages” strategy. Turquoise bars represent the 95% highest posterior density intervals of the ages. Rectangles at the nodes represent the ancestral range with the highest probability based on the inference performed using the younger ages and the genus-level and 10-area coding schemes. When the range encompasses multiple areas, one rectangle is given for each area. Alternative ranges with lower probabilities are provided in Figure S5. P.: Paleocene; EOC.: Eocene, OL.: Oligocene; NEO. + Q.: Neogene + Quaternary.

Ancestral range inferences based on species-level distribution ranges gave similar results (Fig. S6; Table S5), as did the coarser, 7-area coding (Fig. S7; Table S5). The main difference was that the 7-area coding did not result in America being part of the range of ancestral calamoids (Fig. S7). Biogeographic analyses based on the “older ages” resulted in similar range and dispersal inferences as those based on the “younger ages”, except that dispersal events were older (Fig. S8; Table S5). A minimum of 40 dispersals were consistently recovered across the analyses based on the 10-area scheme; their timing based on the “younger ages” is shown in Figure 4. By the end of the Cretaceous, Coryphoideae and Calamoideae had dispersed from northern latitudes to Central America, Arecoideae had dispersed from there to North and South America and to Asia, Coryphoideae and Arecoideae had dispersed to India, Calamoideae and possibly Arecoideae had reached Africa, and Calamoideae had reached localities in Oceania and Europe (Fig. 4A). By the mid-Eocene, Coryphoideae had reached Oceania too and dispersed twice from North America back to Asia. At least six Arecoideae and Coryphoideae dispersals occurred between American regions, and both subfamilies had dispersed recurrently between Asia, India, the Indian Ocean and Madagascar. Arecoideae may also have dispersed to Europe and to Africa during this period (Fig. 4B). Ceroxyloideae are inferred to have dispersed outside Central America by the end of the Oligocene, when they colonised South America, Madagascar and Oceania. Main other dispersals during this period involved Arecoideae, which reached Oceania at least three times and dispersed again to South America, the Indian Ocean and possibly Africa. This is also the first time when Calamoideae are inferred with certainty to have dispersed to South America and from Oceania to Asia, while Coryphoideae reached Africa (Fig. 4C). Dispersal patterns and their consistency between analyses and with the fossil record are further described in Supplementary Results & Discussion based on Supplementary Figures S5-S8.

**Figure 4.**
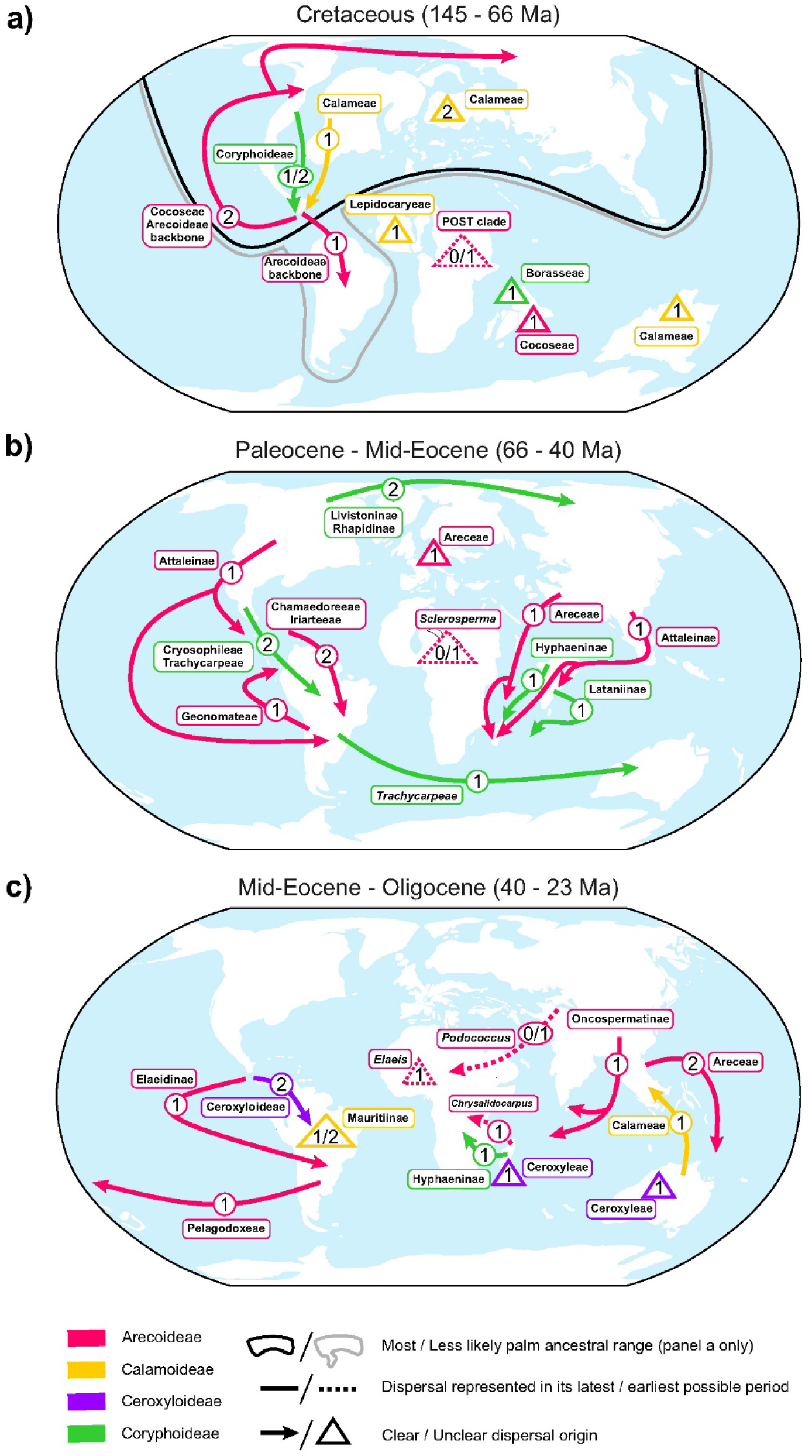
Palm dispersals across the world from the Cretaceous until the Oligocene. **a.** Dispersals inferred to have happened during the Cretaceous, represented on a map of emergent landmasses 92 Ma. **b.** Dispersals inferred to have happened from the Paleocene to mid-Eocene, represented on a map of emergent landmasses 56 Ma. **c.** Dispersals inferred to have happened from the mid-Eocene to the Oligocene, represented on a map of emergent landmasses 34 Ma. POST clade: *Podococcus* + *Orania* + *Sclerosperma* + *Truongsonia*. Numbers indicate the minimum number of dispersals that could be visually inferred based on the ancestral range estimations. Only the dispersals that were congruent across biogeographic inferences based on the 10-area scheme were included. Dispersals are represented in their latest possible period of occurrence based on the “younger ages” results, except for dispersals of Arecoideae in Africa, which are represented (with dashes) in the earliest possible period so that they could be on the figure. When the area of origin of the dispersal was unclear, the dispersal is represented as a triangle in the area of arrival. Maps produced with GPlates 2.5 based on data from Cao et al. (2017) and Müller et al. (2019).

## Discussion

### A Stable Backbone for the Palm Tree Of Life

Here, we provide a fully sampled, genus-level, phylogenetic tree of Arecaceae, supported by evidence from across the nuclear genome. The high level of resolution and support throughout the tree gives us confidence that we have reached a stable understanding of palm relationships (Fig. 1). This new framework broadly agrees with the first comprehensive palm genus phylogenetic tree inferred from 14 plastid and nuclear regions and from morphological data (Baker et al., 2009), but it also delivers insights that can be used to revise the phylogenetic classification of palms. For instance, we provide new evidence for the relationships of genera that were previously unplaced to subtribe in Coryphoideae (unplaced Trachycarpeae genera) and Arecoideae (unplaced Areceae genera; Baker & Dransfield, 2016). We also show that Laccospadicinae should be re-circumscribed to exclude *Calyptrocalyx.* Finally, we identify two non-monophyletic genera (*Oncosperma*, *Areca*), for which systematic solutions must be sought.

Our tree is broadly consistent with the topology for palms recovered in the nuclear phylogenomic, angiosperm-wide tree of Zuntini, Carruthers et al. (2024). Their results are largely based on our Angiosperms353 data, which are re-analysed here in greater depth in concert with PhyloPalm data. The findings of Zuntini, Carruthers et al. (2024) include some anomalies (e.g. Ceroxyloideae embedded in Arecoideae) and are clarified and superseded here by adding more genes and genera. Our study is also in general agreement with the plastome phylogenetic tree of Yao et al. (2023), and provides strongly supported resolution for 58 branches that were weakly supported (and often differently resolved) in the plastome tree, or missing from it. Conversely, the plastome tree shows strongly (though not maximally) supported placements for *Dictyocaryum*, *Sclerosperma*, *Geonoma*, *Acrocomia* and *Leopoldinia*, in disagreement with their different, poorly supported, placements in our nuclear tree (Yao et al., 2023). Both trees therefore complement each other in the elucidation of palm relationships.

### Ancient Gene Flow may have Occurred in Most Palm Subfamilies

Comparison of the plastome tree from Yao et al. (2023) and our nuclear trees reveals areas of the palm tree of life that may have been the theatre of substantial ancient gene flow between lineages. There are ten strongly supported conflicts between the trees, involving Arecoideae (*Wodyetia, Manicaria*, Geonomateae, Iriarteeae), Coryphoideae (*Washingtonia*, *Phoenix*, *Kerriodoxa*) and Calamoideae (*Eugeissona*, *Raphia*, *Calamus*). All conflicts except those involving *Wodyetia* and Iriarteeae are supported by numerous nuclear genes (> 100; Fig. S2). Causes of nuclear-plastome discordance include past hybridisation and incomplete lineage sorting (e.g.: Stull et al., 2020). Most nuclear-plastome conflicts observed here are not associated with ILS/GTE-inconsistent discordance (Figs. 1, S2), which would have been expected in the case of hybridisation. This may indicate incomplete sorting of the plastid genome, or it may still be indicative of hybridisation in a context of “chloroplast capture”, where past hybridisation would have been followed by backcrosses to the paternal parent that erased most of the introgression signal from the nuclear genome, while the maternal plastome was retained (Rieseberg & Soltis, 1991; Tsitrone et al., 2003).

The only cases of nuclear-plastome conflict where ILS/GTE-inconsistent discordance among nuclear genes was observed are those involving *Washingtonia* (and *Phoenix* if not applying the Bonferroni correction), providing further support for the occurrence of past introgression in these taxa, with traces of the maternal genome still detectable in their nuclear genome. The other significant cases of ILS/GTE-inconsistent discordance among nuclear genes (involving *Ammandra*, *Pritchardia* and *Welfia*; Fig. 1) are not associated with nuclear/plastid genome conflicts, suggesting that if there was introgression, the plastome history reflected the majority nuclear history.

### Remaining Polytomies Mainly Result from Rapid Divergences in Arecoideae

A total of 22 branches remain weakly supported despite being underpinned by 86 to 906 nuclear genes (in Cocoseae, Geonomateae, the *Podococcus* + *Orania* + *Sclerosperma* + *Truongsonia* clade and Areceae; Fig. 1). These are likely the result of rapid lineage divergence that prevented the accumulation of sufficient phylogenetic signal and/or the result of widespread discordance among genes due to ILS or other phenomena. All but one of these branches were shorter than the median branch length across dating analyses (Table S4; Fig. S4) and they were not associated with ILS/GTE-inconsistent discordance (Fig. 1), supporting the hypothesis of rapid divergences and suggesting that resolving these polytomies should be possible by comparing more variable regions in lineage-focused studies. The only exception (in Dypsidinae) involved a branch only slightly longer than the median and was associated with ILS/GTE-inconsistent discordance when not applying the Bonferroni correction (Figs. 1, S2; S4), leaving open the possibility that both rapid divergence and hybridisation may contribute to its poor resolution. The latter is further supported by the fact that an in-depth, Dypsidinae-focused, study involving more variable regions also failed to resolve this branch (Eiserhardt et al., 2022). In the case of four other weakly-supported branches, in Coryphoideae (*Pholidocarpus*, Cryosophileae) and Iriarteeae (*Iriartella*), no more than 50 nuclear genes could be assembled (Fig. 1), suggesting that further resolution may easily be attained by producing new, better-quality data.

### Palm Diversification Initiated in the Early Cretaceous and Most Genera Originated by the Oligocene

Our comprehensive sampling and curated fossil records enabled us to provide age estimates for all palm genera and to re-assess previous estimates of palm divergence times. Based on the results from our “younger ages” strategy, we confirm the origin of palms before the end of the Early Cretaceous inferred by Baker & Couvreur (2013a), and we show that most genera (74%) had originated by the Oligocene (Fig. S4). These age estimates tend to be older than in previous studies, and the strongest differences are observed closer to nodes with substantially updated calibrations (e.g. Calamineae and Hyphaeninae), suggesting that discrepancies likely result from the use of older fossils than were available to previous authors. For instance, the median “younger” ages obtained here for the MRCAs of Calamoideae, Ceroxyloideae, Coryphoideae and Arecoideae are all older (by 27–34 Myr) than those found in the latest family-wide study (Baker & Couvreur, 2013a), translating into older ages for tribes and subtribes. Similarly, our median estimates for the palm crown age (119 and 233 Myr; Table S4) were older than those inferred in a recent angiosperm-wide study based on penalised-likelihood molecular dating (90–109 Myr; Zuntini, Carruthers et al., 2024), even though palm stem age estimates overlapped in both studies (134–206 Myr in Zuntini, Carruthers et al., 2024 vs. 120 and 243 Myr here; Table S4). Our age estimates are more consistent with, though still slightly older than those produced in recent studies of palm subgroups (Cano et al., 2022 for American Arecoideae; Escobar et al., 2022 for Ceroxyloideae).

New palm fossils continue to emerge, reminding us of the fragmentary nature of the fossil record and its impact on molecular dating. Our “older ages” analysis acknowledges the incompleteness of the fossil record by permitting older age estimates, which led to the first diversification events in each subfamily being pushed back into the Early Cretaceous. This result receives little support from the fossil record, due to the lack of fossils from this epoch that can be unequivocally attributed to subfamily. Interestingly, the only reliably dated Early Cretaceous palm fossil so far has characters that can be interpreted as ancestral due to their resemblance to both Calamoideae and Nypoideae (Martínez et al., 2016; Table S2). This supports an origin of palms in the Early Cretaceous and their diversification into the current subfamilies mainly taking place in the Late Cretaceous onwards, as recovered with our “younger ages” strategy.

The fossils that we compiled were instrumental to our findings, yet some of them could potentially be placed more precisely in the palm phylogeny. For instance, we confidently assigned only one fossil (corresponding to pollen from the Miocene of Panama; Jaramillo et al., 2014) to Ceroxyloideae, but a few others previously assigned to this subfamily (Brown, 1956; Chate et al., 2019; Kaul, 1943; Khan et al., 2020) have been recently used to calibrate its age (e.g.: Escobar et al., 2022) and may well turn out to be Ceroxyloideae once other possibilities have been ruled out through careful comparative anatomical scrutiny. Targeted paleontological work on palm macro- and micro-fossils is greatly needed to reveal more fossils of interest and unlock the full potential of some fossils already listed for molecular dating and biogeography, notably fossil flowers (Allen, 2015; Chambers et al., 2012; Poinar, 2002b, 2002a; Table S2).

### Palms Dispersed Repeatedly Across Oceanic Barriers

In agreement with the latest family-level biogeographic analyses (Baker & Couvreur, 2013a), we recovered Laurasia as part of the ancestral range of palms, but in addition, we show that the family’s ancestors also occurred in Central America, and possibly South America (Fig. 4). The role of Central America in the early biogeography of palms but also of individual subfamilies (Coryphoideae, Calamoideae) was revealed here by distinguishing more areas than in the previous study. This also enabled us to uncover dispersals to Europe (Calamoideae, Arecoideae) not inferred before. Moreover, although most ancestral ranges and dispersals inferred here were broadly in agreement with those found in Baker & Couvreur (2013a), additional phylogenetic clarity in our study led to a less prominent role of South America during the early history of Arecoideae and Calamoideae, and yielded further insights about how and when Calameae and Pelagodoxeae dispersed to Oceania, Trachycarpeae to Asia, and Ceroxyloideae and Iriarteeae to South America (Fig. 4).

Both studies highlight the important role of long-distance dispersals in the early history of palms, including over large ocean barriers (Fig. 4). For instance, early dispersals of Arecoideae and Coryphoideae to India most likely happened after India was isolated from Africa and before its mid-Eocene collision with Asia (Ali & Aitchison, 2008; Meng et al., 2012), consistent with findings for plants as a whole, as well as reptiles (Klaus et al., 2016). Although palms include some celebrated ocean-dispersed fruits (e.g. coconut, *Nypa*), palm fruits generally do not float and are killed by prolonged salt-water exposure (C. R. Gunn & Dennis, 1976; Kissling, Baker, et al., 2012). Long-distance oceanic dispersal must therefore have involved a vector, such as a bird or floating vegetation mat. These events are considered to be rare, as reflected in the high level of endemism of palm species and higher groups (Kissling, Baker, et al., 2012, Text S1), but our findings indicate that they did occur recurrently over multi-million year timescales (Fig. 4).

Some long-distance dispersals may have been facilitated by plate tectonics creating stepping stones or bringing major landmasses into close proximity. For instance, Central America appears to have served as a stepping stone that enabled Coryphoideae to disperse from North to South America (Fig. 4) long before the Miocene or Pliocene formation of the Isthmus of Panama (O’Dea et al., 2016). Similarly, the dispersals of Trachycarpeae and Trachycarpeae and Pelagodoxeae from America to Oceania in the Paleocene and Eocene also likely happened across large oceanic gaps, but Antarctica was potentially available as a stepping stone, as suggested by the presence of palm pollen and geochemical evidence of a subtropical climate in the area during the Eocene (Korasidis et al., 2022; Pross et al., 2012; Robert & Kennett, 1994). Finally, the narrowing gap between the Sunda and Sahul plates may have facilitated the late Eocene/Oligocene dispersals of Areceae and Calameae between Asia and Oceania well before the Sunda-Sahul collision in the Miocene (Crayn et al., 2015; Hall, 2009), providing new evidence for early initiation of floristic interchange between the plates.

Importantly, we did not infer Late Cretaceous dispersals from Africa to India, nor Eocene dispersals from India to South-East Asia (Africa-India-SE Asia floristic interchange; Morley, 2025), even though extensive fossil evidence supports the existence of such dispersals for *Oncosperma*, *Hydriastele*, *Korthalsia*, *Eugeissona*, Mauritiinae, and Borasseae (Bogotá-Ángel et al., 2021; Morley, 2025; Parmar et al., 2023; Table S2). The reason for this is that relevant fossil records could not be used to inform our ancestral range inferences due to methodological limitations (e.g. node ranges informed by one fossil location, crown nodes of genera not available; Supplementary Methods). This mainly resulted in dispersals from Africa to India and from India to South-East Asia being absent from our reconstructions, but it might also have led to the erroneous inference of an Eocene dispersal of Oncospermatinae from Asia to India, instead of an earlier dispersal in the opposite direction (Fig. 4C). Species-level ancestral range inferences that can incorporate more information from the fossil record will be needed to fully represent the role of Africa and India in the biogeographical history of palms.

### Deep-Time Signatures of Palm Evolution in the Rainforest Biome

Palms have previously been used as a proxy for studying the spatio-temporal origins of the rainforest biome (Couvreur et al., 2011; Couvreur & Baker, 2013). The inference that palms originated 100 Ma in Laurasia in a rainforest-like environment has been used as evidence for the origination of rainforest by the mid-Cretaceous (Couvreur et al., 2011). Our results are broadly consistent with this, notwithstanding older dates and potential involvement of other adjacent areas. Palms then dispersed across the world from the Late Cretaceous until the Paleocene-Eocene Thermal Maximum (56 Ma) while also persisting in Laurasia, which aligns with the occurrence of tropical climates and rainforests in northern latitudes at least intermittently during these times (Burgener et al., 2023; Korasidis et al., 2022). In contrast, from the Oligocene onwards, ancestral ranges only rarely included North America or Europe, and no dispersals to these regions were inferred, consistent with a retraction of the rainforest biome towards equatorial latitudes after the late Eocene-Oligocene global cooling (Scotese et al., 2021; Tardif et al., 2021). Formally testing the hypothesis that palms have been tracking the expansions and retractions of rainforests since their origins will require a much more densely sampled, species-level phylogeny enabling detailed analysis of climatic niche evolution. Overall, our results reflect the climatic constraints inherent in the family (Kissling, Baker, et al., 2012; Reichgelt et al., 2018; Tomlinson et al., 2006) and support the idea that the majority of palm lineages have been megathermal since their origin, hindering their expansion into, and triggering extirpation from, non-megathermal areas in step with climatic changes.

## Conclusions

This study provides a robust phylogenomic and spatio-temporal framework within which to study the evolution of palms. Most relationships among palm genera are now resolved with strong support, potential cases of rapid divergence and reticulation are clearly identified, and needs for taxonomic updates have been highlighted. Our molecular dating and biogeographic analyses showed that palms originated in the Early Cretaceous, that they started to diversify by the end of the Cretaceous, and that palm genera are older than previously estimated. Palms have been dispersing across the world since the Cretaceous, and although these movements may have been facilitated by tectonics at times, the family shows many examples of its ability to cross large distances and oceanic barriers. Accordingly, the pre-Miocene spread of palms across the world was likely mainly determined by their inherent niche conservatism, favouring megathermal areas, in the context of global climatic changes.

Looking forward, clade-specific studies deploying more variable DNA regions and sophisticated coalescent models are required to solve the remaining poorly supported branches and test hypotheses of reticulation. Our fossil review will support focused biogeographic analyses by enabling them to include more fossil information, for instance by using phylogenies including fossil taxa (Heath et al., 2014) as inputs to infer ancestral ranges (Coiro et al., 2023), or by co-estimating ages, fossil placement and ancestral ranges. Performed at the species level, such analyses may shed light on areas mis-represented or missing from the current inferences, such as Africa and India, as well as on the precise effects of climate change on the biogeographic history of the family. Building on the foundations laid here and the ongoing effort towards a species-level phylogenomic tree for palms and other rainforest lineages (Bellot et al., 2020; Murphy et al., 2020), we anticipate an imminent deepening in our understanding of the origins and evolution of the tropical rainforest biome (Eiserhardt et al., 2017).

## Supplementary Material

Supplementary Methods, Supplementary Results & Discussion, Supplementary Figures and Supplementary Tables are available from GitHub at: https://github.com/sidonieB/Bellot_et_al_Palm_Early_Evolution_Supplementary_Material together with the raw and clean sequence alignments, the gene trees and the species trees generated for this study. (The final version of the supplementary material will be deposited in a Dryad Digital Repository once the study has been peer-reviewed and accepted for publication).

## Supporting information

Supplementary Methods, Supplementary Results & Discussion and Supplementary Figures

## Acknowledgements

We thank Rodrigo Bernal, Grace Brewer, Boris Domenech, John Dowe, Niroshini Epitawalage, Isabel Fairlie, Sophie Nadot, Turkan Ozdemir, and staff from SING Herbarium, Fairchild Tropical Gardens and the Naturalis Biodiversity Center for providing or helping generate data from up to five samples, and Laszlo Csiba for general support in the lab. We are grateful to Lisa Pokorny for discussions and support in early phases of the project and for sharing data from three samples.We acknowledge the Research/Scientific Computing teams at The James Hutton Institute and NIAB for providing computational resources and technical support for the “UK’s Crop Diversity Bioinformatics HPC” (BBSRC grant BB/S019669/1), use of which has contributed to the results reported within this paper.

## Authors contributions

S. B.: Conceptualization, Methodology, Software, Validation, Formal analysis, Investigation, Resources, Data Curation, Writing – original draft, Writing – Review & Editing, Visualization, Supervision; F. L. C: Methodology, Software, Validation, Formal analysis, Investigation, Resources, Writing – Review & Editing, Visualization. K. K. S. M.: Conceptualization, Investigation, Data Curation, Writing - Review & Editing; R. J. M: Investigation, Data Curation, Writing - Review & Editing; A. C.: Ressources, Data Curation, Writing – Review & Editing; T. L. C.: Conceptualization, Ressources, Writing – Review & Editing; R. C.: Supervision; W. L. E.: Conceptualization, Writing – Review & Editing; B. G. K.: Investigation, Writing – Review & Editing; O. M.: Data Curation, Writing – Review & Editing; M. S.: Investigation; F. F.: Supervision, Writing – Review & Editing, Funding acquisition; I. J. L.: Supervision, Funding acquisition; W. J. B: Conceptualization, Methodology, Validation, Investigation, Resources, Data Curation, Writing – original draft, Writing – Review & Editing, Visualization, Supervision, Project administration, Funding acquisition.

