## Supplementary Methods, Supplementary Results & Discussion and Supplementary Figures for "Early Cretaceous Origin and Evolutionary History of Palms (Arecaceae) inferred from 1,033 Nuclear Genes and a New Synthesis of Fossil Evidence"

The supplementary material provided below is available from GitHub at: https://github.com/sidonieB/Bellot_et_al_Palm_Early_Evolution_Supplementary_Material, together with the raw and clean sequence alignments, the gene trees and the species trees generated for this study.

The final version of the supplementary material will be deposited in a Dryad Digital Repository once the study has been peer-reviewed and accepted for publication.

**Contents:**

**Supplementary Methods** (provided below)
**Supplementary Results & Discussion** (provided below)
**Supplementary Tables** (provided in separate excel workbook Bellot_et_al_Origin_and_Evolution_of_Palms_Supplementary_Tables_S1_S2_S3_S4_S5.xlsx) including:

- **Supplementary Table S1. Sampling and biogeographical area coding**
- **Supplementary Table S2. Palm fossils**
- **Supplementary Table S3. Summary of molecular dating analyses**
- **Supplementary Table S4. Molecular dating results**
- **Supplementary Table S5. Biogeographical analyses results**

**Supplementary Figures** (provided below) including:

**Supplementary Figure S1. Consistency of median age estimates across molecular dating analyses.** For both strategies (“younger ages” and “older ages”), median age estimates are compared between analyses including (7 calibrations) or excluding (6 calibrations) *F. messelensis* as part of the calibration points, and between analyses constraining or not the tree topology to reflect that of the species tree inferred with ASTRAL.

**Supplementary Figure S2. Phylogenetic relationships among palm genera. Left:** all-genes tree based on 802 nuclear genes classified as paralogous and 231 classified as orthologous; **Right:** orthologs-only tree based only on the 231 genes classified as orthologous. Labels above branches indicate local posterior probabilities. Stars indicate branches associated with ILS/GTE-inconsistent discordance without (*) or with (**) Bonferroni correction. Numbers below branches indicate the number of gene trees in which that branch was recovered. Pie charts represent quartet scores, with black representing the scores of the displayed topology while grey and white represent the scores of the alternative topologies. Red arrows indicate strongly supported conflicts between the all-genes tree and the plastome phylogenetic tree inferred by Yao et al. (2023).

**Supplementary Figure S3. Relationship between gene number, branch resolution and gene tree discordance.** The classification of branches associated or not with ILS/GTE-inconsistent discordance was performed with (top) and without (bottom) Bonferroni correction.

**Supplementary Figure S4. Variation in divergence time estimates between the "younger ages" and "older ages" strategies.** Median ages (bold numbers) and 95% highest posterior density intervals (bars) were obtained by applying the older ages (left) and the younger ages (right) strategies to a constrained topology without using *F. messelensis* as a calibration point. Small numbers correspond to clades listed in Table S4. The inset shows the relationship between branch length (calculated as the difference between the median ages of the parent and child nodes of the branch) and support in the all-genes species tree shown in Figure 1 (Strong: local posterior probability $\geq 0.9$; Weak: local posterior probability $< 0.9$) after excluding four branches that were supported by no more than 50 genes.

**Supplementary Figure S5. Ancestral ranges estimated based on the younger ages and genus-level tip ranges coded following the 10-area scheme.** Rectangles at the nodes represent the ancestral range with the highest probability – when the range encompasses multiple areas, multiple rectangles are printed on the same line. The second and third most probable ranges are printed above and under the nodes for cases where the most probable range had a probability inferior to 0.66 or to twice the probability of the second most probable range. Pie charts indicate the probability of the most probable range (black), second most probable range (grey), third most probable range (light grey) and of all other ranges cumulated (white).

**Supplementary Figure S6. Ancestral ranges estimated based on the younger ages and species-level tip ranges coded following the 10-area scheme.** Rectangles at the nodes represent the ancestral range with the highest probability – when the range encompasses multiple areas, multiple rectangles are printed on the same line. The second and third most probable ranges are printed above and under the nodes for cases where the most probable range had a probability inferior to 0.66 or to twice the probability of the second most probable range. Pie charts indicate the probability of the most probable range (black), second most probable range (grey), third most probable range (light grey) and of all other ranges cumulated (white).

**Supplementary Figure S7. Ancestral ranges estimated based on the younger ages and genus-level tip ranges coded following the 7-area scheme.** Rectangles at the nodes represent the ancestral range with the highest probability – when the range encompasses multiple areas, multiple rectangles are printed on the same line. The second and third most probable ranges are printed above and under the nodes for cases where the most probable range had a probability inferior to 0.66 or to twice the probability of the second most probable range. Pie charts indicate the probability of the most probable range (black), second most probable range (grey), third most probable range (light grey) and of all other ranges cumulated (white).

**Supplementary Figure S8. Ancestral ranges estimated based on the older ages and genus-level tip ranges coded following the 10-area scheme.** Rectangles at the nodes represent the ancestral range with the highest probability – when the range encompasses multiple areas, multiple rectangles are printed on the same line. The second and third most probable ranges are printed above and under the nodes for cases where the most probable range had a probability inferior to 0.66 or to twice the probability of the second most probable range. Pie charts indicate the probability of the most probable range (black), second most probable range (grey), third most probable range (light grey) and of all other ranges cumulated (white).

**Supplementary Methods**

*Tissue sampling and DNA sequencing*

All currently recognised palm genera were represented by one species. Additional species were sampled in two genera (*Areca*, *Oncosperma*) that may not be monophyletic, based on preliminary analyses. The genus *Dasypogon* (Dasypogonaceae) was included as an outgroup, following most recent monocot phylogenomic studies (Barrett et al., 2016; Li et al., 2021). Plant material was obtained from silica-dried leaf tissue or from herbarium specimens. Species names, sample origin and voucher information are provided in Table S1.

After DNA extraction, DNA longer than ca. 1,000 base pairs (bp) was fragmented using a M220 Focused-ultrasonicator™ and AFA Fiber Pre-Slit Snap-Cap microTUBES (Covaris, Woburn, MA, USA), with the following settings: peak power: 50; duty % factor: 20; cycles/burst: 200; power: 10; duration: 55 seconds; temperature: 20°C. Hybridisation lasted 24 hours and was conducted separately for the Angiosperms353 and PhyloPalm probe kits (Johnson et al., 2019; Loiseau et al., 2019), with a temperature of hybridisation of 65°C. It was followed by 12–16 PCR cycles as recommended in the manufacturer's protocol (http://www.arborbiosci.com/mybaits-manual). Libraries enriched in Angiosperms353 or PhyloPalm regions were pooled and sequenced separately, generating two sequence datasets for each sample, except for 23 samples (out of 187) for which only PhyloPalm data could be generated, and 2 for which only Angiosperms353 data could be generated, because there was not enough DNA library to perform two hybridisations (the type of data that was generated for each sample is indicated in Table S1). DNA sequencing was performed on an Illumina MiSeq with v2 or v3 chemistry at the Royal Botanic Gardens, Kew, or on an Illumina HiSeq X at Macrogen Inc. (Seoul, Korea), yielding 2 x 300 bp-long or 2 x 150 bp-long paired-end reads, respectively. For nine taxa (Table S1), the PhyloPalm dataset had been generated prior to this study and was downloaded from public repositories or provided by co-authors. The raw sequencing read data was deposited in GenBank (accession numbers in Table S1). Parts of the data that were newly generated as part of this study were used in two recent studies: Angiosperm353 regions from most genera were included in a study on the diversification of all angiosperms (Zuntini, Carruthers et al., 2024) and a small subset of the PhyloPalm regions (known as the Heyduk regions; Heyduk et al., 2016) from 48 samples was used in studies of the palms of New Caledonia (Pérez-Calle, Bellot et al., 2024) and Madagascar (Eiserhardt et al., 2022).

*Genomic data cleaning and assembly*

Sequence data quality was assessed using FASTQC v. 0.11.9 (Andrews, 2010), and Trimmomatic v. 0.39 (Bolger et al., 2014) was used to remove Illumina sequencing adapters using the paired-end palindrome mode with 1 seed mismatch allowed, palindrome and simple clips thresholds of 30 and 7 respectively, a minimum adapter length of 2 bp and keeping both reads. Bases with low quality at the end of the reads were then removed using a sliding window ("SLIDINGWINDOW" parameter) of 4 bp and a minimum Phred quality threshold of 30, while bases of low quality at the beginning of the reads were removed using the

“LEADING” parameter and the same minimum quality threshold. Reads shorter than 40 bp after the trimming were discarded.

Clean reads were analysed using HybPiper v. 1.3.1 (Johnson et al., 2016) to recover and assemble the regions targeted by both probe kits. The original reference sequences used to recover reads matching the target regions comprised the default Angiosperms353 reference sequences (https://github.com/mossmatters/Angiosperms353/Angiosperms353_targetSequences.fasta), and the sequences used to design the PhyloPalm probe kit, mostly derived from the *Elaeis guineensis* genome (K. Heyduk, com. pers.; M. Paris, com. pers.; Heyduk et al., 2016; Loiseau et al., 2019). Using blastn 2.5.0 (part of the BLAST+ suite; Camacho et al., 2009), we found that 68 Angiosperms353 target regions aligned with the PhyloPalm target regions over more than 50 bp, with an e-value < 0.05, so the Angiosperms353 duplicated regions were discarded from the original reference sequences so that the same region would not be recovered and analysed twice. To maximise gene recovery across all samples, a custom set of reference sequences was built that included all the original reference sequences as well as target sequences recovered by submitting five high-quality datasets to an initial HybPiper run using the original reference sequences. The five datasets comprised data from species representing the five palm subfamilies: *Attalea butyracea* (Arecoideae), *Korthalsia echinometra* (Calamoideae), *Phytelephas aequatorialis* (Ceroxyloideae), *Chuniophoenix nana* (Coryphoideae) and *Nypa fruticans* (Nypoideae). This final set of reference sequences comprises 11,348 sequences corresponding to 1,255 target regions and is available at https://github.com/sidonieB/Palm_phylogenomics_resources/blob/main/A353PP_noOverlap_285-970_wPalmRefs.fasta.

*Classification of paralogous and orthologous regions*

Previous studies have provided evidence for the occurrence of a whole genome duplication event in the ancestral lineage that gave rise to all current palm species (Barrett et al., 2019), suggesting that some or all the genetic regions targeted by our study may originally have been duplicated in the ancestral lineage, even if they now appear to be single copy regions. This issue has previously been addressed for subfamily Calamoideae by using published whole genome sequences of *Calamus* (Zhao et al., 2018) to identify genes with multiple copies. To mitigate for the risk of aligning and analysing paralogs instead of orthologs across the palm family, we expanded an approach first developed for Calamoideae (Kuhnhäuser, 2021) by classifying the regions into orthologous vs. paralogous regions based on their copy numbers in the annotated genomes of palm species representing the three main palm subfamilies Arecoideae, Calamoideae and Coryphoideae: *Elaeis guineensis* (Arecoideae; coding sequences from the EG5 genome version 3 downloaded from http://genomsawit.mpob.gov.my on 20 November 2020; Chan et al., 2017), *Calamus simplicifolius* (Calamoideae; coding sequences file downloaded from http://gigadb.org/dataset/101052 on 8 March 2020; Zhao et al., 2018) and *Phoenix dactylifera* (Coryphoideae; transcript variants downloaded from https://datepalmgenomehub.abudhabi.nyu.edu/?q=node/12; Hazzouri et al., 2019). Our target regions were aligned against all the coding sequences from the annotated genome and a region was considered paralogous if the search identified at least two hits with the coding sequence, each with e-value scores < $10^{-21}$ and overlapping by more than 30% of the coding sequence length. This was conducted separately for each of the three annotated genomes, and a region identified as paralogous based on at least one of these searches was considered paralogous, while regions not identified as paralogous based on any of the three searches were considered orthologous. The latter is based on the rationale that even if these regions were duplicated in the ancestral lineage that gave rise to palms, the fact that they are found to be single copy regions in calamoids, coryphoids and arecoids suggests that reduction to a single copy occurred before these subfamilies diverged from each-other, and therefore before palms diversified, leading to the same copy being subsequently shared by all subfamilies and species. In addition, we classified as paralogous any region that raised a "paralog warning" in at least one sample during the HybPiper analysis. These warnings are raised when multiple overlapping, different contigs are assembled from the reads matching a single target region, which can indicate that the region is present in multiple copies in the sample being analysed. Classifying such regions as paralogous therefore allowed us to mitigate the risk of paralogy due to more recent genome/gene duplications that might have taken place throughout palm evolution. This paralog search resulted in classifying 951 out of 1,255 target regions as paralogous and 304 as orthologous

*Cleaning of multiple sequence alignments*

Multiple sequence alignments were generated for each region using MAFFT v. 7 (Katoh & Standley, 2013) with the "--genafpair" setting and 1,000 iterations. OptrimAl (https://github.com/keblat/bioinfo-utils/blob/master/docs/advice/scripts/optrimAl.txt) was then used to trim alignment columns so as to minimise gaps in alignments while keeping their informativeness high. Trimmed alignments were then further cleaned with CIAlign v. 1.1.0 (Tumescheit et al., 2022) to remove sequences that were very divergent from the others and therefore likely spurious. After trying different thresholds, sequences were removed if less than 85% of their nucleotides corresponded to the most common base in the alignment at that position. Care was taken to not discard the outgroup taxon sequence data during this process by using the option "--retain-str". TAPER v. 1.0 (Zhang et al., 2021) was then used on the resulting alignments to remove mis-aligned, highly divergent stretches of otherwise non-spurious sequences, using default settings. The resulting alignments were then trimmed again with OptrimAl. As a result of the cleaning process, 137 target regions were discarded because they contained less than four taxa or because the alignment entirely comprised sequences shorter than 250 bp or considered uninformative by OptrimAl.

*Molecular dating analyses*

Times of divergence between palm genera were estimated using BEAST 2.6 (Bouckaert et al., 2019). Because of computational limitations and potential conflicts between gene histories, we based the molecular dating analysis on the 22 orthologous regions that yielded the top 10% gene trees with the highest bipartition agreement to the species tree based on orthologous genes, as identified by (Smith et al., 2018). These 22 regions were concatenated using AMAS (Borowiec, 2016) and BEAST was run allowing each region to be analysed under a different nucleotide substitution model corresponding to the one identified previously using IQ-TREE v. 1.6.12 (Minh et al., 2020), or to the closest model available in

BEAST. Clocks and trees were linked across the partitions to enable the analyses to converge in a reasonable time. Substitution rate priors were set to lognormal distributions to facilitate chain mixing, and the tree generation prior was set according to the incomplete sampling birth-death model. To account for substitution rate heterogeneity between taxa, we used an optimised relaxed clock (ORC) model (Douglas et al., 2021). Analyses (see next section) were performed with and without constraining the tree topology based on the ASTRAL species tree obtained from summarizing all the genes. Topological constraints were implemented by modifying some parameters in the BEAST xml files following recommendations at https://www.beast2.org/2014/07/28/all-about-starting-trees and https://www.beast2.org/fix-starting-tree/. Specifically, we specified the species tree topology and we set the weights of the subtreeslide, narrowExchange, wideExchang, wilsonBalding and ORCAdaptableOperatorSampler_NER parameters to 0 directly in the xml configuration files. We used Tracer v. 1.7.2 (Rambaut et al., 2018) to monitor the convergence of the parameter estimates and to check that effective sampling sizes were mostly above 200. A run without using the data ("prior-only") was also performed for each analysis, in order to verify that the prior settings did not drive the posterior parameter estimations more than the data themselves. Number of runs, generations and effective sampling sizes are provided in Table S3 for each analysis. A maximum clade credibility tree was generated from the posterior tree distribution of each analysis using TreeAnnotator, which is part of BEAST, after removing the first 10% posterior trees generated (burn-in fraction). Median ages estimated for each clade across the post-burn-in posterior tree distribution were reported on this tree, together with the 95% highest posterior density age intervals.

*Fossil calibrations for the molecular dating*

We used fossil records to calibrate the age of six nodes spread across the palm family. *Mauritiidites* spp. pollen records from the Maastrichtian to Campanian of Nigeria, Sudan, Egypt, Cameroon, Gabon and Angola (Edet & Nyong, 1993; Eisawi & Schrank, 2008; Rull, 1998; Salami, 1990; Salard-Cheboldaeff, 1990; Schrank, 1994; Table S2) were used to calibrate the age of the stem node of Mauritiinae. Pollen records of *Dicolpopollis* spp. from the Cenomanian (Burger, 1990; Macphail & Jordan, 2015; Totterdell & Mitchell, 2009) were used to calibrate the stem node of the clade made of Metroxylinae, Plectocomiinae, Calaminae and Pigafettinae (Calamoideae, Calameae). These pollen grains have thin exine, which could support a more internal placement excluding Metroxylinae, but we refrain from doing this until further studies confirm this placement (Table S2). *Sabal bigbendense* Manch., Wheeler, & Lehman from the Campanian of Texas (ca. 77 Ma; (Cano, 2018; Manchester et al., 2010) was used to calibrate the stem node of *Sabal* (Coryphoideae, Sabaleae). *Hyphaenocarpon indicum* Bande, Prakash, & Ambwani emend. Matsunaga, S.Y.Sm., Manch., Srivastava, & Kapgate (Bande et al., 1982; Matsunaga et al., 2019) from the Deccan Intertrappean Beds (64–67 Ma; Matsunaga et al., 2018) was used to calibrate the crown node of Hyphaeninae (Coryphoideae, Borasseae). *Palmocarpon drypetoides* (Mehrotra, Prakash & Bande) Manch., Bonde, Nipunage, Srivastava, Mehrotra & S.Y.Sm. (Manchester et al., 2016), also from the Deccan Intertrappean Beds (64 – 67 Ma; Matsunaga et al., 2018), was used to calibrate the crown of Attaleinae (Arecoideae, Cocoseae). Finally, the fossil *Friedemannia messelensis* Collinson, Manch. & Wilde from the middle Eocene of Germany (ca. 47 Ma; Collinson et al., 2012; Matsunaga & Smith, 2021) was used to calibrate the crown of Areceae (Arecoideae). The placement of these last three fossils was supported by previous morphology-based phylogenetic inference (Matsunaga & Smith, 2021). Further details on the fossils are provided in Table S2.

*Ancestral range inferences using DEC*

The evolution of ancestral ranges along the palm phylogeny was estimated using the Dispersal-Extinction-Cladogenesis (DEC) model (Ree & Sanmartín, 2018), as implemented in the DEC eXtended version (DECX, Beeravolu & Condamine, 2016; available at: https://github.com/champost/DECX). DECX requires a time-calibrated tree, the current distribution of each taxon for a set of geographic areas, and a time-stratified geographic model that is represented by connectivity and dispersal multiplier matrices for specified time intervals spanning the entire evolutionary history of the group. DECX allows classical vicariance as a cladogenetic event by using temporally flexible constraints on the connectivity between any two given areas following the movement of landmasses and dispersal opportunity over time. However, DECX does not incorporate the founder-event speciation (+J parameter) because of concerns with statistical validity of model choice among DEC-derived models (Ree & Sanmartín, 2018). Also, founder-event speciation often leads to inferences that are decoupled from time, with null or extremely low extinction rates, an effect of the model favouring cladogenetic events over anagenetic events (Ree & Sanmartín, 2018), which makes it inadequate for reconstructing the history of ancient groups with widespread distributions.

*Area delimitation for the ancestral range inferences*

Analyses were performed using two area delimitation schemes. The first scheme (10-area) comprised North America, Central America, South America, Europe, Asia, Africa, Oceania, India, Indian Ocean, and Madagascar. The second scheme (7-area) comprised North America (including Central America from the 10-area scheme) South America, Africa, India, Eurasia (merging Europe and Asia from the 10-area scheme), Oceania and Indian Ocean (including Madagascar from the 10-area scheme). The delimitation between Asia and Oceania was different in each scheme (following the Lidekker line in the 10-area scheme, and the Wallace line in the 7-area scheme; see map below). Using both schemes enabled us to explore the impact of different rationales subtending area definitions, and to compare results to the main previous global palm biogeography study (Baker & Couvreur, 2013) that used the 7-area scheme.

**Maps of both area schemes:**

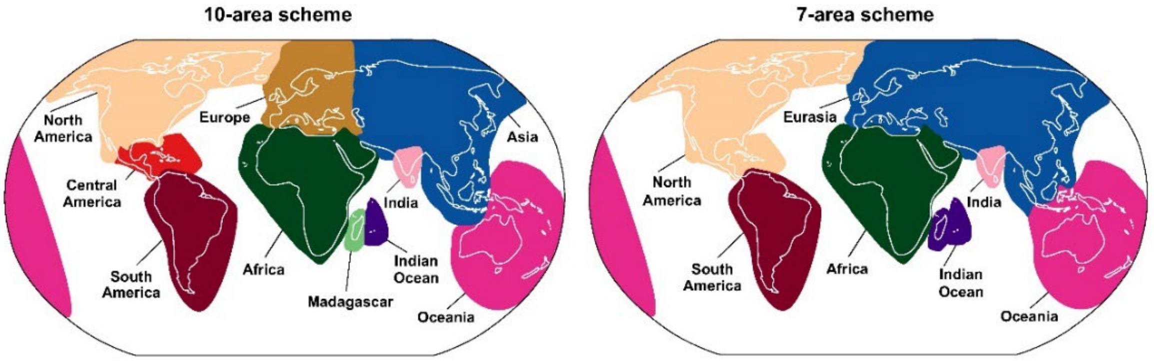


The table on the following page provides the precise definition and underlying rationale for each area for each scheme.

| 10-area scheme | Delimitation | Rationale (based on geology during periods relevant for genera divergences, i. e. Oligocene and older) |
|---|---|---|
| North America | Includes Florida, Bermuda, and northern Mexico down to Sinaloa, Durango, Zacatecas, Nuevo Leon, San Luis Potosi and Tamaulipas included | The rest of Mexico was not formed/emerged as early as the part included here |
| Central America | Antilles (but not Trinidad), and everything between Panama and the Mexico states excluded from North America | Trinidad is geologically part of South America. Central America as defined here had a geological history very different from North and South America. |
| South America | Includes Trinidad and everything south of Central America | See Central America |
| Europe | From the British isles until the Turgai strait | The Turgai strait was the main potential barrier across Eurasia |
| Asia | From the Turgai strait to everything north west of Lydekker line, including Wallacea, Nicobar island, Andaman islands. | In the past, Wallacea was physically closer to South East Asia than to New Guinea |
| Africa | Africa, the Arabian Peninsula, Canary Islands, Cape Verde | The Arabian Peninsula was closer to Africa than Asia for a long part of their geological history |
| Oceania | Includes Australia, New Guinea, New Zealand, and everything south east of Lydekker line, including the south Pacific islands | See Asia. Assignment of south Pacific islands to Oceania was decided because a detailed study of the spread of palms across the Pacific can only be meaningfully done at the species level and by delimiting many new areas corresponding to each archipelago. |
| India | Includes Sri Lanka, Assam, and Bengladesh | These regions were either not emerged or part of India when it was not yet attached to Asia |
| Indian Ocean | Mascarenes, Seychelles | Fragments of areas which were much bigger in the past, and which were separated from India and Madagascar |
| Madagascar | Includes Comoros | Assignment of Comoros to Madagascar instead of Africa is arbitrary, they could be on their own in a more precise analysis but they concern very few palms so we chose to avoid adding an area just for these. |
| **7-area scheme** | | |
| North America | Includes everything north of Panama (included) + Antilles (but not Trinidad) | Follows Baker & Couvreur 2013; Merges North America and Central America of Scheme 1 |
| South America | Includes everything south of Panama, incl. Trinidad | Follows Baker & Couvreur 2013 |
| Eurasia | From the British Isles to Wallace's line, including Andaman and Nicobar islands | Follows Baker & Couvreur 2013; Merges Europe and a slightly reduced Asia from Scheme 1 (excludes Wallacea) |
| Oceania | Everything East of Wallace's line, incl. Australia, New Guinea, New Zealand, and the Pacific islands | Follows Baker & Couvreur 2013; slightly expanded compared to Scheme 1 (includes Wallacea) |
| Africa | Includes the Arabian Peninsula, Canary Islands, Cape Verde | Follows Baker & Couvreur 2013 |
| India | Includes Sri Lanka, Assam, Bengladesh | Follows Baker & Couvreur 2013 |
| Indian Ocean | Mascarenes, Seychelles, Madagascar, Comoros | Follows Baker & Couvreur 2013; Merges Indian Ocean and Madagascar from Scheme 1 |

*Connectivity information for the ancestral range inferences*

Analyses were performed under a model where area connectivities were constrained based on how distances between areas changed through geological periods, after preliminary analyses showed that analyses performed without such constraints provided unrealistic results where ancestral species would be spread across many distant areas. Based on paleogeographic reconstructions (e.g.: Kocsis & Scotese, 2021; Seton et al., 2012), we built connectivity matrices to represent major changes in tectonic conditions that could have affected the distribution of palms. Constraints on area connectivity were specified in a matrix by coding 0 if any two areas are not connected or 1 if these areas are connected during a given period. The time interval was dissected into four time slices: from 0 to 36 Ma, 36 to 56 Ma, 56 to 85 Ma, and 85 Ma to the tree root age. Hence, a time-stratified geographic model was created in the form of binary matrices that consider paleogeographic changes through time with time slices indicating the possibility or not for a species to colonize a new area (Beeravolu & Condamine, 2016).

Antarctica was not included in the area schemes as no palm currently occurs there and no palm fossil has been found there so far, which means that nothing in the data could directly indicate the occurrence of a palm lineage in this area. However, an occurrence in Antarctica remains implicitly allowed by the model via the connectivity matrices allocating non-null probabilities to connections between Oceania, Africa and America during the Cretaceous, when these regions were connected through what is now Antarctica.

The tables on the following page show the connectivity matrices for each scheme. In these matrices, a ‘1’ at the intersection of two areas means that a taxon can occupy a range encompassing the two areas, while a ‘0’ means that it cannot occupy a range encompassing the two areas.

**10-area scheme**

| **200 - 86 Ma** | **NA** | **CA** | **SA** | **EU** | **AF** | **IN** | **AS** | **OC** | **IO** | **MA** | **56 - 37 Ma** | **NA** | **CA** | **SA** | **EU** | **AF** | **IN** | **AS** | **OC** | **IO** | **MA** |
|---|---|---|---|---|---|---|---|---|---|---|---|---|---|---|---|---|---|---|---|---|---|
| **North America** | 1 | | | | | | | | | | **North America** | 1 | | | | | | | | | |
| **Central America** | 1 | 1 | | | | | | | | | **Central America** | 1 | 1 | | | | | | | | |
| **South America** | 1 | 1 | 1 | | | | | | | | **South America** | 0 | 1 | 1 | | | | | | | |
| **Europe** | 1 | 1 | 1 | 1 | | | | | | | **Europe** | 1 | 0 | 0 | 1 | | | | | | |
| **Africa** | 1 | 1 | 1 | 1 | 1 | | | | | | **Africa** | 0 | 0 | 0 | 1 | 1 | | | | | |
| **India** | 1 | 1 | 1 | 0 | 1 | 1 | | | | | **India** | 0 | 0 | 0 | 0 | 0 | 1 | | | | |
| **Asia** | 1 | 1 | 1 | 1 | 0 | 0 | 1 | | | | **Asia** | 1 | 0 | 0 | 1 | 0 | 1 | 1 | | | |
| **Oceania** | 1 | 1 | 1 | 0 | 1 | 1 | 0 | 1 | | | **Oceania** | 0 | 0 | 1 | 0 | 0 | 0 | 0 | 1 | | |
| **Indian Ocean** | 1 | 1 | 1 | 0 | 1 | 1 | 0 | 1 | 1 | | **Indian Ocean** | 0 | 0 | 0 | 0 | 1 | 1 | 1 | 0 | 1 | |
| **Madagascar** | 1 | 1 | 1 | 0 | 1 | 1 | 0 | 1 | 1 | 1 | **Madagascar** | 0 | 0 | 0 | 0 | 1 | 1 | 0 | 0 | 1 | 1 |
| **85 - 57 Ma** | | | | | | | | | | | **36 - 0 Ma** | | | | | | | | | | |
| **North America** | 1 | | | | | | | | | | **North America** | 1 | | | | | | | | | |
| **Central America** | 1 | 1 | | | | | | | | | **Central America** | 1 | 1 | | | | | | | | |
| **South America** | 0 | 1 | 1 | | | | | | | | **South America** | 1 | 1 | 1 | | | | | | | |
| **Europe** | 1 | 0 | 0 | 1 | | | | | | | **Europe** | 1 | 0 | 0 | 1 | | | | | | |
| **Africa** | 0 | 0 | 0 | 1 | 1 | | | | | | **Africa** | 0 | 0 | 0 | 1 | 1 | | | | | |
| **India** | 0 | 0 | 0 | 0 | 1 | 1 | | | | | **India** | 0 | 0 | 0 | 1 | 1 | 1 | | | | |
| **Asia** | 1 | 0 | 0 | 1 | 0 | 0 | 1 | | | | **Asia** | 1 | 0 | 0 | 1 | 1 | 1 | 1 | | | |
| **Oceania** | 0 | 0 | 1 | 0 | 0 | 0 | 0 | 1 | | | **Oceania** | 0 | 0 | 0 | 0 | 0 | 0 | 1 | 1 | | |
| **Indian Ocean** | 0 | 0 | 0 | 0 | 1 | 1 | 0 | 0 | 1 | | **Indian Ocean** | 0 | 0 | 0 | 0 | 1 | 1 | 1 | 0 | 1 | |
| **Madagascar** | 0 | 0 | 0 | 0 | 1 | 1 | 0 | 0 | 1 | 1 | **Madagascar** | 0 | 0 | 0 | 0 | 1 | 1 | 0 | 0 | 1 | 1 |

**7-area scheme**

| **200 - 86 Ma** | **NA** | **SA** | **AF** | **IN** | **EU** | **OC** | **IO** | **56 - 37 Ma** | **NA** | **SA** | **AF** | **IN** | **EU** | **OC** | **IO** |
|---|---|---|---|---|---|---|---|---|---|---|---|---|---|---|---|
| **North America** | 1 | | | | | | | **North America** | 1 | | | | | | |
| **South America** | 1 | 1 | | | | | | **South America** | 0 | 1 | | | | | |
| **Africa** | 1 | 1 | 1 | | | | | **Africa** | 0 | 0 | 1 | | | | |
| **India** | 1 | 1 | 1 | 1 | | | | **India** | 0 | 0 | 0 | 1 | | | |
| **Eurasia** | 1 | 1 | 1 | 0 | 1 | | | **Eurasia** | 1 | 0 | 1 | 1 | 1 | | |
| **Oceania** | 1 | 1 | 1 | 1 | 0 | 1 | | **Oceania** | 0 | 1 | 0 | 0 | 1 | 1 | |
| **Indian Ocean** | 1 | 1 | 1 | 1 | 0 | 1 | 1 | **Indian Ocean** | 0 | 0 | 1 | 1 | 1 | 0 | 1 |
| **85 - 57 Ma** | | | | | | | | **36 - 0 Ma** | | | | | | | |
| **North America** | 1 | | | | | | | **North America** | 1 | | | | | | |
| **South America** | 0 | 1 | | | | | | **South America** | 1 | 1 | | | | | |
| **Africa** | 0 | 0 | 1 | | | | | **Africa** | 0 | 0 | 1 | | | | |
| **India** | 0 | 0 | 1 | 1 | | | | **India** | 0 | 0 | 1 | 1 | | | |
| **Eurasia** | 1 | 0 | 1 | 0 | 1 | | | **Eurasia** | 1 | 0 | 1 | 1 | 1 | | |
| **Oceania** | 0 | 1 | 0 | 0 | 0 | 1 | | **Oceania** | 0 | 0 | 0 | 0 | 1 | 1 | |
| **Indian Ocean** | 0 | 0 | 1 | 1 | 0 | 0 | 1 | **Indian Ocean** | 0 | 0 | 1 | 1 | 1 | 0 | 1 |

*Fossil selection for ancestral range inferences*

We explored the possibility of informing the ancestral distribution ranges of some clades based on the information available in the palm fossil record (Table S2). Fossil assignment is provided at the clade level, including or not the stem of the clade. For instance, a fossil assigned to Coryphoideae stem + crown means that we know that the fossil belongs to a taxon more closely related to Coryphoideae as we know them today than to any other palm, which means that the taxon may have been the "stem" lineage directly leading to extant coryphoids, or it could have been a lineage deriving from this stem but now extinct, or it could have been a lineage from inside the crown of coryphoids, again either a direct ancestor of an extant coryphoid or a now-extinct derived coryphoid (see stars in figure below). If this fossil occurred in, let's say, Africa, this information would be fed to the DEC implementation used here by constraining the range of the crown node of Coryphoideae to include at least Africa (without penalising other areas). By doing this, the model would allow transitions to ranges including Africa at the node itself, but also along the stem leading to that node, and Africa would become a more likely area to be included in the ranges of more derived nodes (see figure below). For this reason, fossil choice was restricted to fossils that could be assigned to internal nodes, while fossils assigned to tip branches could not be exploited as it would require recoding the tip distribution to include the fossil range (see Figure below).

Among fossils that could be used, we only selected the fossils that had an age range overlapping with the 95% highest posterior density age interval of the branch leading to the node they were assigned to. This means that the set of fossils selected was allowed to change depending on the input dated tree.

When multiple fossils satisfying these criteria were available for the same node, we selected the fossil coming from an area that was not represented in the distribution range of the extant species included in the clade to which the fossil was assigned. If multiple fossils satisfied this criterion, we selected the oldest one. This enabled to maximise the informativeness of the selected fossil set. Because ages were different between the "younger ages" and "older ages" trees, the analysis using the former tree could be informed by eight fossils while the latter could be informed by four of these eight fossils. The selected fossils included some used to calibrate the molecular dating as well as some other fossils. They are highlighted in Table S2, which also explains for each unselected fossil, why it was not selected.

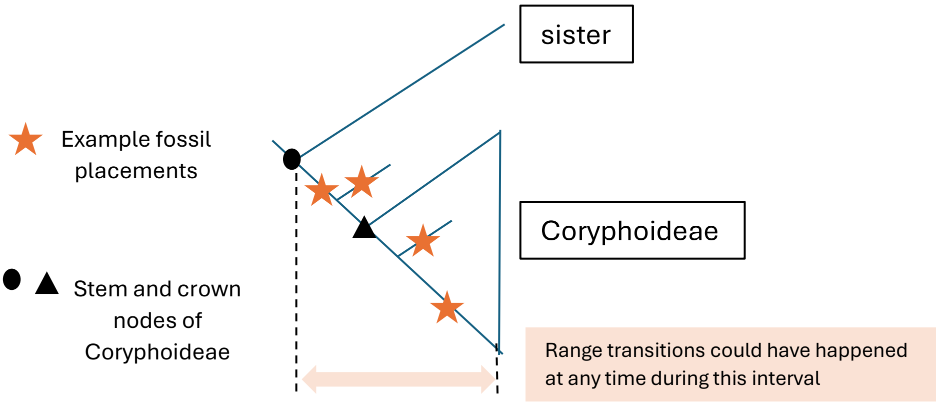


**Fossil selection explanatory figure**. In this example, an African fossil is known to belong to Coryphoideae (without knowing if belonging to the stem or crown) and we assume that Coryphoideae do not occur in Africa today. Assigning the fossil to the crown node (triangle) constrains the range of that node to include Africa, and allows transitions to Africa to have occurred along the stem of Coryphoideae. In contrast, assigning the fossil to the stem node would have constrained the range of the stem node to include Africa and therefore allowed transitions to Africa along the stem leading to both Coryphoideae and their sister, even though there is no data suggesting that the latter may have had ancestors in Africa. For this reason, fossils assigned to the stem + crown of a lineage (i.e. most fossils) were assigned to the crown node of that lineage in the context of the DECX analysis. Following this reasoning, fossils assigned to tips (in this study genera or species) but occurring in different areas than the corresponding extant genus/species could not be used because it would be wrong to code the extant taxon as occurring in an area where it does not occur anymore, and because assigning the fossil to the stem node of the tip would mean assuming that the transition in area happened before the tip separated from its sister.

**Supplementary Results & Discussion**

*Phylogenetic relationships in each subfamily*

In Calamoideae, relationships among Lepidocaryeae showed Raphiinae strongly supported as sister to Ancistrophyllinae. Eugeissoneae were strongly supported as sister to Calameae, in which Korthalsiinae, Salaccineae, Metroxylinae and Pigafettinae formed a strongly supported grade around sister clades Calaminae and Plectocomiinae (Fig. 1). The two latter subtribes were resolved as sisters in both trees, with strong support in the all-genes tree (Fig. S2). There was no ILS/GTE-inconsistent discordance in this subfamily.

In Coryphoideae, the "syncarpous clade" (Borasseae, Caryoteae, Corypheae, Chuniophoeniceae; Dransfield et al., 2008) was strongly supported as sister to the "CSPT clade" (Cryosophileae, Sabaleae, Phoeniceae, Trachycarpeae; Baker & Dransfield, 2016). Tribes Caryoteae and Chuniophoeniceae formed a grade around sister tribes Corypheae and Borasseae, with all relationships in the clade being identically resolved and strongly supported in both trees (Fig. 1). Sabaleae and Cryosophileae were strongly supported as sisters. In Cryosophileae, *Trithrinax* and *Leucothrinax* had different placements in both trees, with weak support in at least the orthologs-only tree (Fig. S2). Other relationships in the tribe were identically resolved in both trees. Phoeniceae was strongly supported as sister to the Sabaleae-Cryosophileae clade in the all-genes tree, but weakly supported as sister to these and Trachycarpeae in the orthologs-only tree (Fig. S2, inset on Fig. 1). This conflicting placement of Phoeniceae was significantly associated with ILS/GTE-inconsistent discordance, but only when the Bonferroni correction was not applied to the test. In Trachycarpeae, *Brahea* (unplaced at the subtribe level in the current classification; Baker & Dransfield, 2016) was strongly supported as sister to Rhapidinae; *Serenoa* and *Acoelorrhaphe* (both unplaced) were strongly supported as a clade sister to Livistoninae. Relationships in Rhapidinae and Livistoninae were identically resolved in both trees. *Colpothrinax* (previously unplaced) was strongly supported as sister to all Trachycarpeae except *Pritchardia*, *Copernicia* and *Washingtonia* (all previously unplaced). The relationships among these three genera and the remaining of Trachycarpeae were associated with ILS/GTE-inconsistent discordance (Fig. 1).

In Ceroxyloideae, Cyclospatheae was resolved as sister to Phytelepheae. All relationships were identically resolved in both trees, but ILS/GTE-inconsistent discordance was found within Phytelepheae.

In Arecoideae, Iriarteeae, Chamaedoreeae, the "POST clade" (*sensu* Baker et al., 2011, modified by Sâm et al., 2023; Podococceae, Oranieae, Sclerospermeae, Truongsonieae) and the "RRC clade" (*sensu* Baker et al., 2011; Reinhardtieae, Roystoneae, Cocoseae) formed a strongly supported grade around the "core arecoid clade" (*sensu* Baker et al., 2011; Areceae, Euterpeae, Geonomateae, Leopoldinieae, Manicarieae, Pelagodoxeae). Relationships in Iriarteeae were identically resolved in both trees (mainly with strong support) and those in Chamaedoreeae were all strongly supported and identically resolved in both trees. The POST clade comprises two clades, one with Podococceae strongly supported as sister to Truongsonieae, the other (weakly supported in the orthologs-only tree) made of Oranieae and Sclerospermeae (Figs. 1 and S2). In the RRC clade, which is sister to the core arecoid clade, Cocoseae was strongly resolved as sister to Reinhardtieae, and both were strongly supported as sister to Roystoneae. In Cocoseae, most branches were identically resolved and strongly supported in both trees (except for the placements of *Syagrus* and *Cocos*, weakly supported in the orthologs-only tree). However, relationships among Bactridinae differed between both trees (and had weak support in at least the orthologs-only tree; Fig. S2).

In the core arecoid clade, Euterpeae were strongly resolved as sister to Areceae. and both were strongly resolved as most closely related to Leopoldinieae, Manicarieae, Pelagodoxeae and Geonomateae than to Cocoseae + Reinhardtieae + Roystoneae. Geonomateae, Manicarieae and Pelagodoxeae formed a strongly supported clade in which the latter two were sisters in both trees. The placement of Leopoldinieae was different and weakly supported in both trees (Fig. S2). In Geonomateae, the position of *Geonoma* was resolved differently depending on the tree, in both cases with weak support (Fig. S2), while other relationships were all strongly supported and identically resolved in both trees. However, there was ILS/GTE-inconsistent discordance about *Welfia*'s placement in the all-genes dataset (Fig. 1). In Euterpeae, all relationships were strongly supported and identically resolved in both trees.

Among Areceae, the "west Pacific clade" (Baker et al., 2011; comprising Archontophoenicinae, Basseliniinae, Carpoxylinae, Clinospermatinae, Laccospadicinae, Ptychospermatinae, Rhopalostylidinae, *Dransfieldia* and *Heterospathe*) was strongly supported, and sister to *Hydriastele* with strong support in both trees. Inside the west Pacific clade, *Calyptrocalyx* was excluded from Laccospadicinae with strong support, resolving as sister to Archontophoenicinae, and Rhopalostylidinae formed a clade nested in Basseliniinae, although the placement is weakly supported. Most branches in the west Pacific clade were identically resolved in both trees and strongly supported in at least one of the trees. Three exceptions were *Veillonia*, *Lepidorrhachis* and *Howea*, the placements of which were differently resolved depending on the tree, with weak support in both trees (Fig. S2). The conflicting placement of *Veillonia* was associated with ILS/GTE-inconsistent discordance when the Bonferroni correction was not applied (Fig. 1). Outside the west Pacific clade, *Clinostigma*, *Cyrtostachys* and *Bentinckia* (unplaced at the subtribe level; Baker & Dransfield, 2016) formed a strongly supported grade in which Arecinae is nested. The unplaced *Rhopaloblaste* and *Dictyosperma* were strongly supported as sister genera in both trees. Relationships in Arecinae, Oncospermatinae, Verschaffeltiinae and Dypsidinae were all identically resolved in both trees, mostly with strong support. The species *Areca chaiana* and *Oncosperma fasciculatum*, which we added to our sampling because we suspected that they may render their genera non-monophyletic, were each strongly supported as placing outside of their respective genera. Relationships among subtribes outside the west Pacific clade and the exact placements of *Iguanura*, *Bentinckia* and *Loxococcus* were resolved differently in both trees, always with weak support (Figs. 1 and S2). This was not associated with ILS/GTE-inconsistent discordance.

*Ancestral ranges and dispersals in each subfamily*

All inferences suggested that the MRCA of palms occurred in Laurasia and Central America (Fig. 3, Figs. S5-S8), and possibly also in South America when accounting for alternative states with similar probabilities (Fig. S5). The latter is in agreement with the discovery of a palm pollen fossil in Patagonia (Argentina) from the Albian that bears characters that can be interpreted as ancestral to the family due to their resemblance to both

Calamoideae and Nypoideae (Martínez et al., 2016; Table S2). The uncertainty around the ancestral range of palms and the areas of origin of the earliest dispersals (Fig. 4) may remain until the oldest fossils can be more reliably placed in the family so that their geographic distribution can inform biogeographical inferences. Meanwhile, an early, if not ancestral, presence of the family throughout Laurasia remains highly credible given the existence of palm fossils from the Turonian of Europe (Crié, 1892; Kvaček & Herman, 2004), the Santonian/Coniacian of North America (Berry, 1916; Greenwood et al., 2022) and the Campanian of Asia (Harley, 2006; Takahashi, 1964).

The ancestral range of Calamoideae most likely comprised North America and Asia, and possibly already Europe and Central America, where the subfamily was present by the end of the Cretaceous. By then, Calamoideae had also reached Africa (MRCA of Lepidocaryeae), Europe again (Salaccinae stem node, younger ages only) and Oceania (stem node of *Metroxylon*), and they may have dispersed back from Asia to North and Central America (Calameae), although alternative states at this node support instead a continuous presence in those regions (Figs. 3, S5). Mauritiinae appear to have dispersed twice to South America, at least once by the mid-Eocene-Oligocene (Figs. S5-S8), or possibly the Paleocene when considering age confidence intervals (Fig. S4), in agreement with the existence of Maurittinae fossils from the paleocene of South America (Bogotá-Ángel et al., 2021). Around the mid-Eocene-Oligocene, a dispersal from Oceania to Asia occurred in Calameae, possibly in the stem lineage of Plectocomiinae (Figs. 3, S5). The 7-area coding did not result in America being recovered as part of the range of ancestral calamoids (Fig. S7). Given that the oldest known Calamoideae fossils from Oceania have been assigned to internal nodes of the subfamily, the fact that the most ancestral ranges of Calamoideae do not include Oceania is not in direct contradiction with the fossil record. However, it is surprising that so far, no Cretaceous Calamoideae fossils have been recorded from the areas inferred here to have been part of their ancestral range (North and Central America, Europe, Asia) - this requires further investigation, as illustrated by the high uncertainty surrounding range estimates at the deepest nodes of this subfamily (Fig. S5).

The ancestors of Coryphoideae most likely occurred in Asia and North America (Figs. 3, S5). By the end of the Cretaceous, the subfamily had dispersed once or twice to Central America, and once to India (Borasseae stem). The arrival of the subfamily to South America most likely involved two independent dispersals from Central/North America (in Cryosophileae and the MRCA of *Copernicia* + *Pritchardia* + *Washingtonia*) during the Paleocene or Eocene, although a single more ancestral or to the contrary even more later arrivals were inferred when using the older ages (Fig. S8) or the second area coding (Fig. S7). Coryphoideae dispersed from India to the Indian Ocean and Madagascar (Borasseae), from America to Oceania (*Pritchardia-Copernicia* stem) and twice from North America to Asia (Livistoninae and Rhapidineae stems) at the latest by the mid-Eocene (Figs 3, S5). Finally, they reached Africa when the MRCA of *Hyphaene*+*Medemia* dispersed there from India or most likely Madagascar, by the end of the Eocene (Fig. S5).

The most likely range of Ceroxyloideae's MRCA was Central America, possibly together with North America (Fig. S5). The presence of the subfamily in South America was inferred to be either the result of two independent dispersals that happened in Phytelepheae and Ceroxyleae by the Oligocene (Figs. 3, S5, S6, S7), or of a single ancestral dispersal (older ages; Fig. S8). Ceroxyleae dispersed from America to Oceania at the latest by the end of the Oligocene and then to Madagascar (*Ravenea*), possibly involving Asia and/or India as stepping stones (younger ages only; Figs. S5, S6, S7).

Finally, the MRCA of Arecoideae was inferred to have occurred at least in Central America, and possibly also in North and South America and Asia (Figs. 3, S5-S8). The subfamily was present in all these regions and in India by the end of the Cretaceous (backbone nodes, Cocoseae; Figs. 3, S5-S8). Since then, Arecoideae have been dispersing again multiple times between the American regions (Chamaedoreeae, Iriarteeae, Geonomateae, Attaleinae, Elaeidinae; Figs. 3, S5). The subfamily had dispersed at least once to the Indian Ocean region and twice to Madagascar by the mid-Eocene (along the stems of Attaleinae and Areceae), and once again to the Indian Ocean by the end of this period (Oncospermatinae), with these dispersals most likely involving Asia and India as areas of origin (Figs. 3, S5). The ancestors of Pelagodoxeae had dispersed from South America to Oceania by the end of the Oligocene, and possibly much earlier (Figs. S5-S8). The arrival of Areceae (west Pacific clade) in Oceania was inferred to have occurred at the same period via two dispersals from Asia (younger ages; Figs. 3, S5, S6), although it could have happened as early as the Late Cretaceous via a single dispersal from America or Asia (older ages or second area scheme; Figs. S8, S7). At least four independent dispersals of Arecoideae to Africa occurred: in *Podococcus*, in *Sclerosperma* (or in the POST clade MRCA), in *Jubaeopsis* (or the Attaleineae MRCA), in *Elaeis* and in *Chrysalidocarpus* (Figs 3, S5-S8). Even when only considering younger ages, these dispersals were difficult to date due to the long branches involved, and could have occurred from the Late Cretaceous (POST clade MRCA), the Paleocene (*Sclerosperma*), the Eocene (*Jubaeopsis*, *Podococcus*) or the Oligocene (*Elaeis*, *Chrysalidocarpus*) to the Quaternary (Figs. 3, S5). The presence of ancestral Areceae in Europe was inferred when using younger ages, most likely resulting from a dispersal from Asia or America (Figs. 3, S5). In contrast, results based on older ages did not recover the presence of Areceae in Europe, but instead suggested that Attaleinae ancestors may have occurred there (Fig. S8).

Figure S1

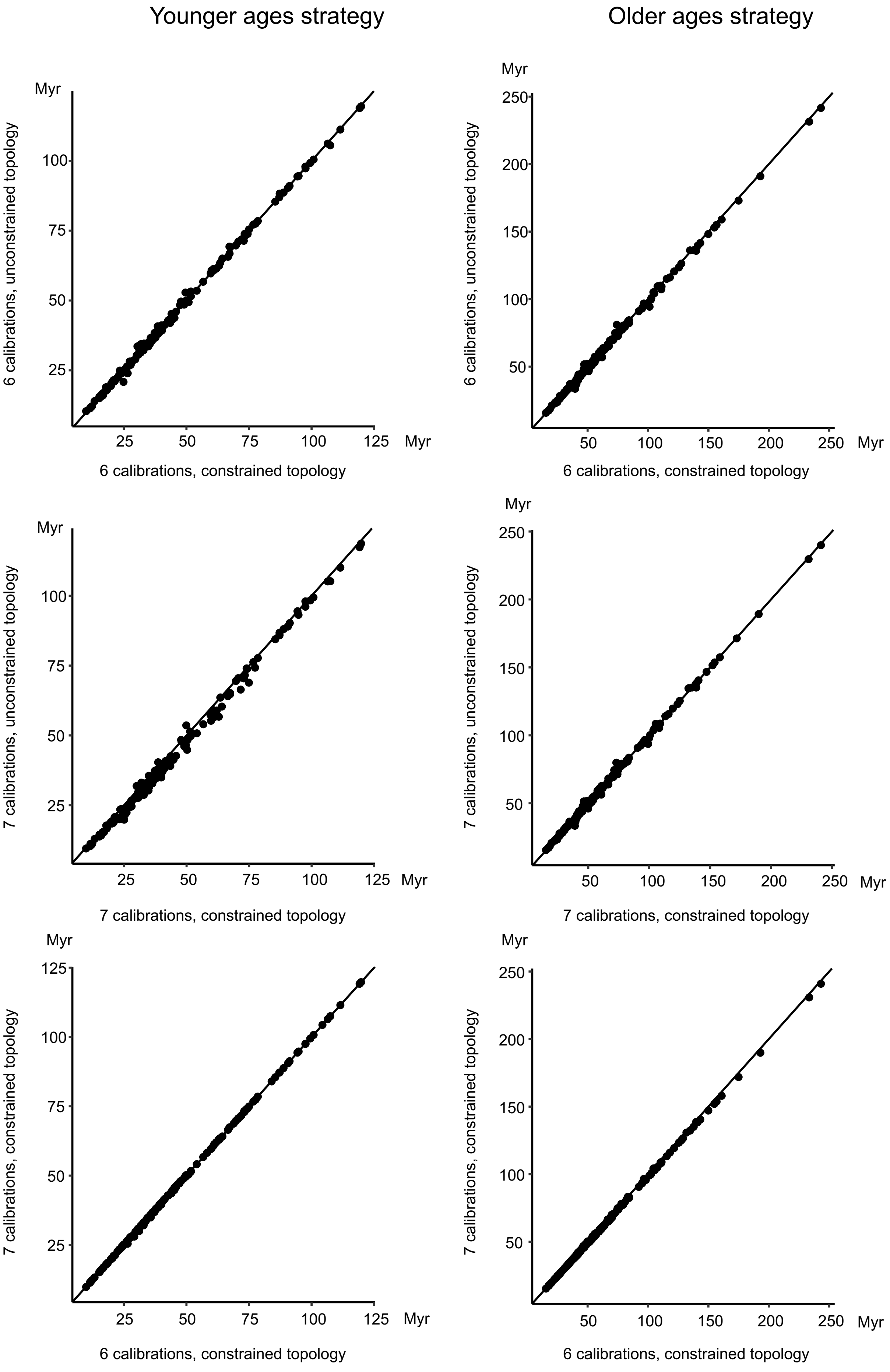
Younger ages strategy
Older ages strategy
Myr
6 calibrations, unconstrained topology
6 calibrations, constrained topology
7 calibrations, unconstrained topology
7 calibrations, constrained topology
25
50
75
100
125
150
200
250

Figure S2

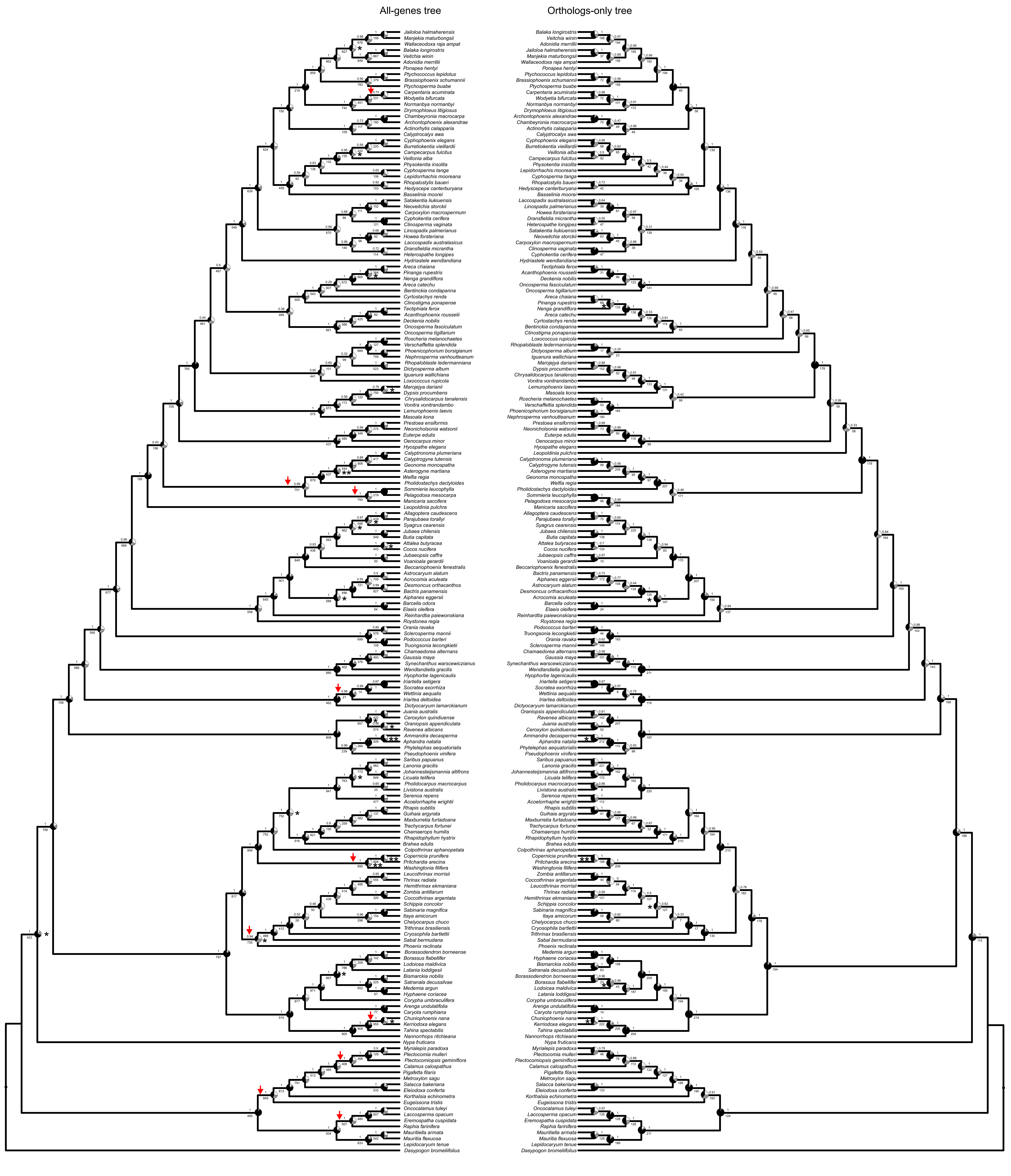

Figure S3

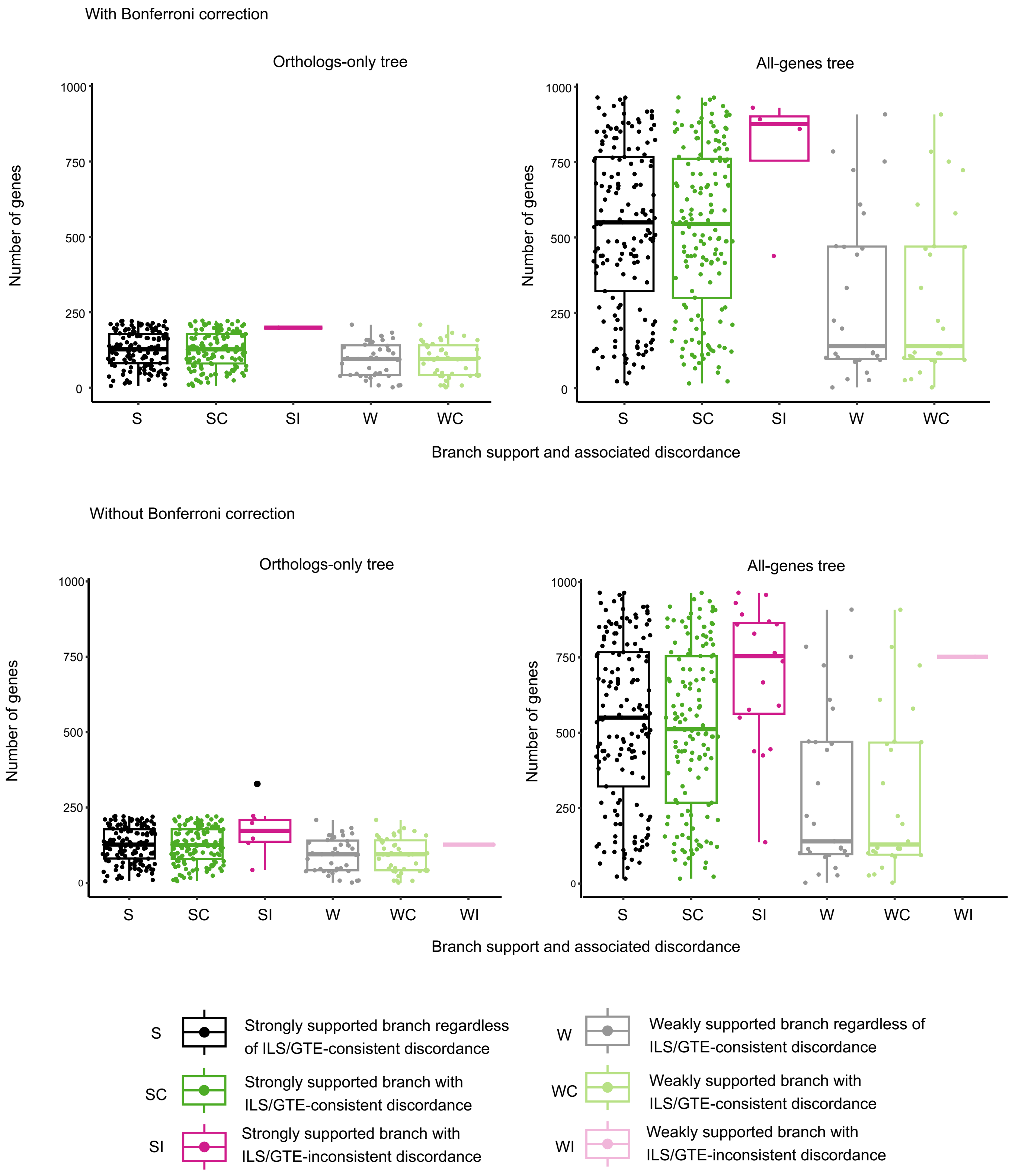
With Bonferroni correction
Orthologs-only tree
All-genes tree
Number of genes
0
250
500
750
1000
S
SC
SI
W
WC
Branch support and associated discordance
Without Bonferroni correction
Orthologs-only tree
All-genes tree
WI
S
Strongly supported branch regardless of ILS/GTE-consistent discordance
SC
Strongly supported branch with ILS/GTE-consistent discordance
SI
Strongly supported branch with ILS/GTE-inconsistent discordance
W
Weakly supported branch regardless of ILS/GTE-consistent discordance
WC
Weakly supported branch with ILS/GTE-consistent discordance
WI
Weakly supported branch with ILS/GTE-inconsistent discordance

Figure S4

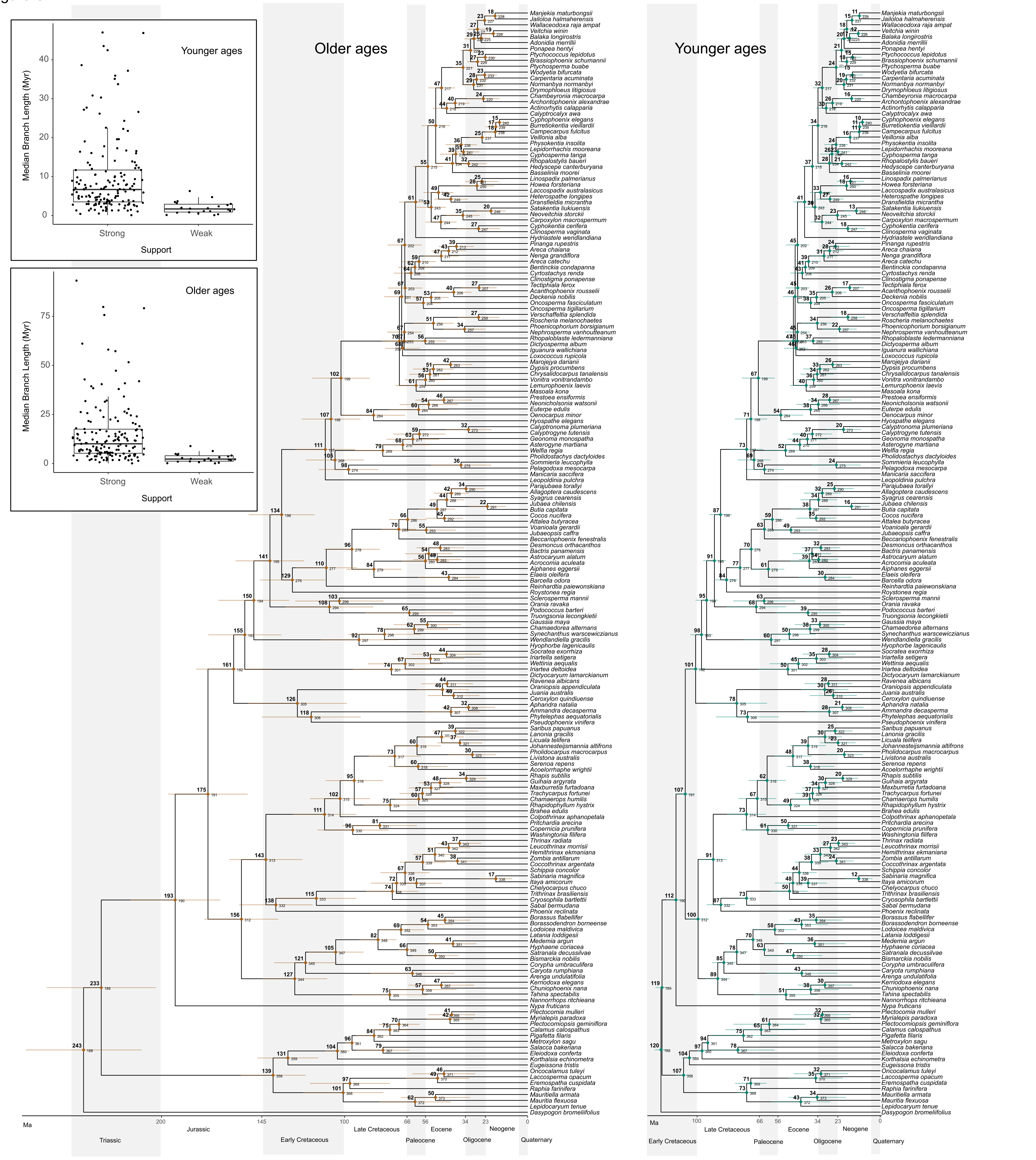
Younger ages
Median Branch Length (Myr)
Strong
Weak
Support
Older ages
Median Branch Length (Myr)
Strong
Weak
Support
Older ages
Younger ages
Ma
Triassic
Jurassic
Early Cretaceous
Late Cretaceous
Paleocene
Eocene
Oligocene
Neogene
Quaternary

Figure S5

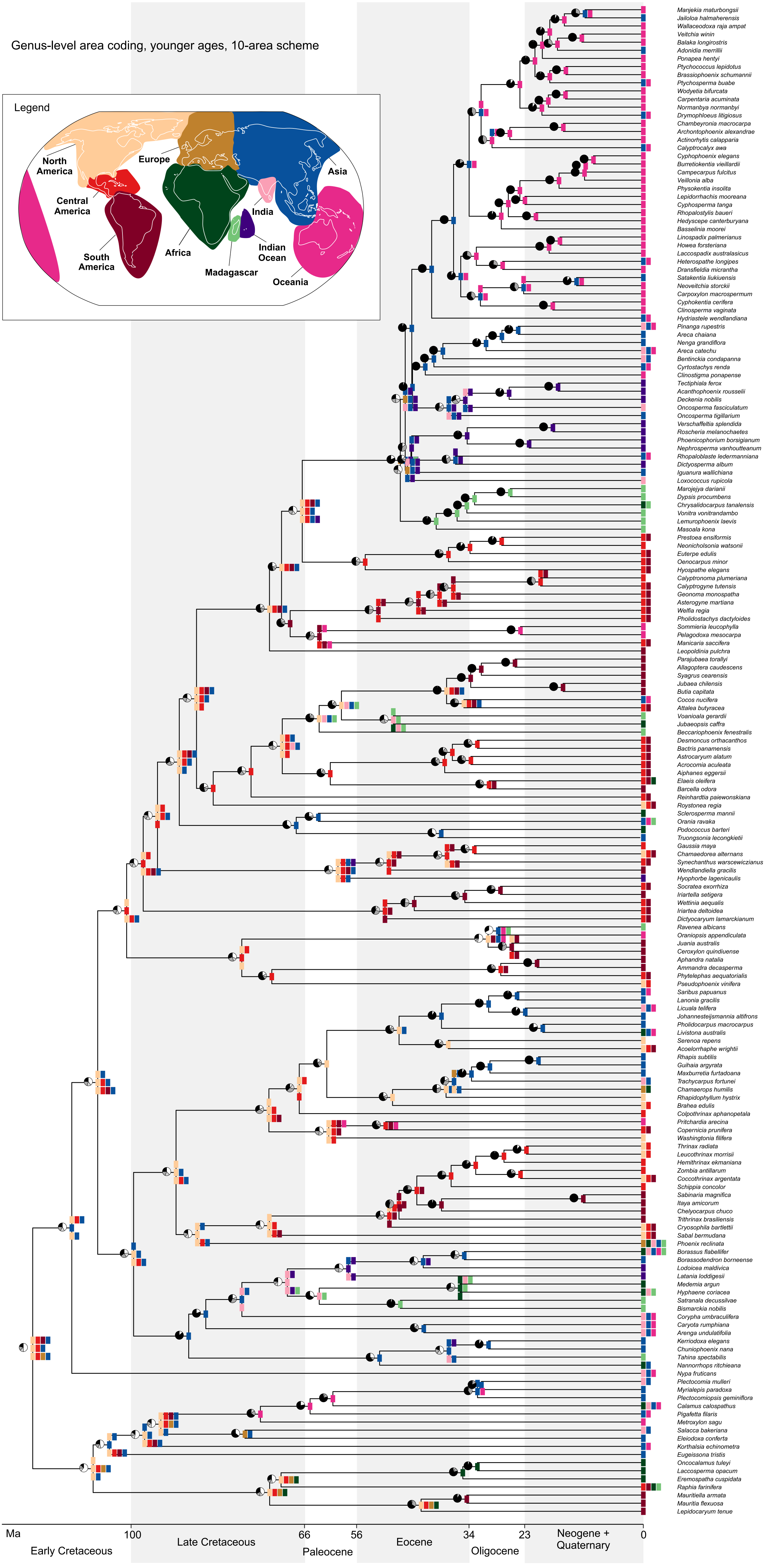
Genus-level area coding, younger ages, 10-area scheme
Legend
North America
Europe
Asia
Central America
India
South America
Africa
Indian Ocean
Madagascar
Oceania
Manjekia maturbongsii
Jailoloa halmaherensis
Wallaceodoxa raja ampat
Veitchia winin
Balaka longirostris
Adonidia merrillii
Ponapea hentyi
Ptychococcus lepidotus
Brassiophoenix schumannii
Ptychosperma buabe
Wodyetia bifurcata
Carpentaria acuminata
Normanbya normanbyi
Drymophloeus litigiosus
Chambeyronia macrocarpa
Archontophoenix alexandrae
Actinorhytis calapparia
Calyptrocalyx awa
Cyphophoenix elegans
Burretiokentia vieillardii
Campecarpus fulcitus
Veillonia alba
Physokentia insolita
Lepidorrhachis mooreana
Cyphosperma tanga
Rhopalostylis baueri
Hedyscepe canterburyana
Basselinia moorei
Linospadix palmerianus
Howea forsteriana
Laccospadix australasicus
Heterospathe longipes
Dransfieldia micrantha
Satakentia liukiuensis
Neoveitchia storckii
Carpoxylon macrospermum
Cyphokentia cerifera
Clinosperma vaginata
Hydriastele wendlandiana
Pinanga rupestris
Areca chaiana
Nenga grandiflora
Areca catechu
Bentinckia condapanna
Cyrtostachys renda
Clinostigma ponapense
Tectiphiala ferox
Acanthophoenix rousselii
Deckenia nobilis
Oncosperma fasciculatum
Oncosperma tigillarium
Verschaffeltia splendida
Roscheria melanochaetes
Phoenicophorium borsigianum
Nephrosperma vanhoutteanum
Rhopaloblaste ledermanniana
Dictyosperma album
Iguanura wallichiana
Loxococcus rupicola
Marojejya darianii
Dypsis procumbens
Chrysalidocarpus tanalensis
Vonitra vonitrandambo
Lemurophoenix laevis
Masoala kona
Prestoea ensiformis
Neonicholsonia watsonii
Euterpe edulis
Oenocarpus minor
Hyospathe elegans
Calyptronoma plumeriana
Calyptrogyne tutensis
Geonoma monospatha
Asterogyne martiana
Welfia regia
Pholidostachys dactyloides
Sommieria leucophylla
Pelagodoxa mesocarpa
Manicaria saccifera
Leopoldinia pulchra
Parajubaea torallyi
Allagoptera caudescens
Syagrus cearensis
Jubaea chilensis
Butia capitata
Cocos nucifera
Attalea butyracea
Voanioala gerardii
Jubaeopsis caffra
Beccariophoenix fenestralis
Desmoncus orthacanthos
Bactris panamensis
Astrocaryum alatum
Acrocomia aculeata
Aiphanes eggersii
Elaeis oleifera
Barcella odora
Reinhardtia paiewonskiana
Roystonea regia
Sclerosperma mannii
Orania ravaka
Podococcus barteri
Truongsonia lecongkietii
Gaussia maya
Chamaedorea alternans
Synechanthus warscewiczianus
Wendlandiella gracilis
Hyophorbe lagenicaulis
Socratea exorrhiza
Iriartella setigera
Wettinia aequalis
Iriartea deltoidea
Dictyocaryum lamarckianum
Ravenea albicans
Oraniopsis appendiculata
Juania australis
Ceroxylon quindiuense
Aphandra natalia
Ammandra decasperma
Phytelephas aequatorialis
Pseudophoenix vinifera
Saribus papuanus
Lanonia gracilis
Licuala telifera
Johannesteijsmannia altifrons
Pholidocarpus macrocarpus
Livistona australis
Serenoa repens
Acoelorrhaphe wrightii
Rhapis subtilis
Guihaia argyrata
Maxburretia furtadoana
Trachycarpus fortunei
Chamaerops humilis
Rhapidophyllum hystrix
Brahea edulis
Colpothrinax aphanopetala
Pritchardia arecina
Copernicia prunifera
Washingtonia filifera
Thrinax radiata
Leucothrinax morrisii
Hemithrinax ekmaniana
Zombia antillarum
Coccothrinax argentata
Schippia concolor
Sabinaria magnifica
Itaya amicorum
Chelyocarpus chuco
Trithrinax brasiliensis
Cryosophila bartlettii
Sabal bermudana
Phoenix reclinata
Borassus flabellifer
Borassodendron borneense
Lodoicea maldivica
Latania loddigesii
Medemia argun
Hyphaene coriacea
Satranala decussilvae
Bismarckia nobilis
Corypha umbraculifera
Caryota rumphiana
Arenga undulatifolia
Kerriodoxa elegans
Chuniophoenix nana
Tahina spectabilis
Nannorrhops ritchieana
Nypa fruticans
Plectocomia mulleri
Myrialepis paradoxa
Plectocomiopsis geminiflora
Calamus calospathus
Pigafetta filaris
Metroxylon sagu
Salacca bakeriana
Eleiodoxa conferta
Korthalsia echinometra
Eugeissona tristis
Oncocalamus tuleyi
Laccosperma opacum
Eremospatha cuspidata
Raphia farinifera
Mauritiella armata
Mauritia flexuosa
Lepidocaryum tenue
Ma
100
66
56
34
23
0
Early Cretaceous
Late Cretaceous
Paleocene
Eocene
Oligocene
Neogene + Quaternary

Figure S6

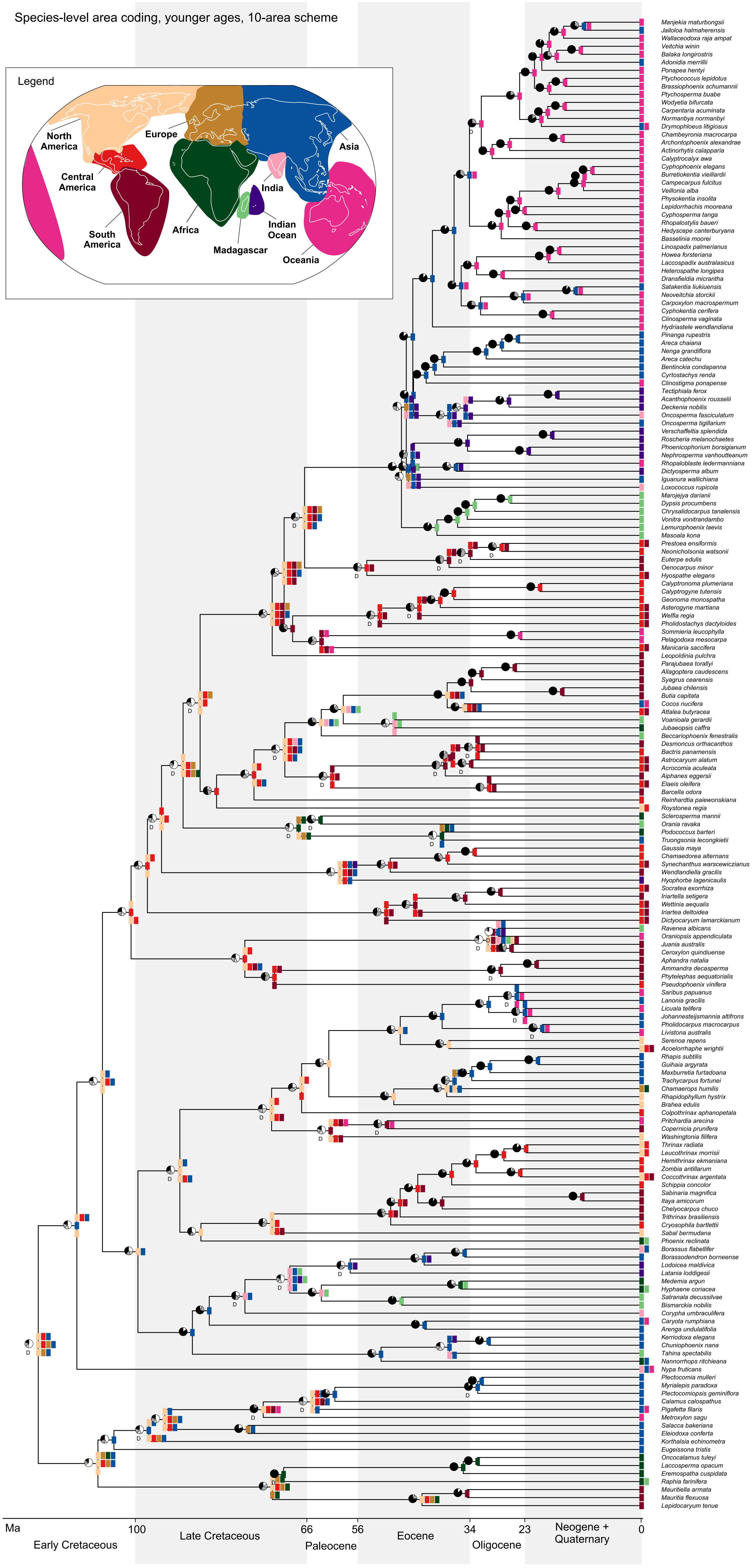
Species-level area coding, younger ages, 10-area scheme
Legend
North America
Europe
Asia
Central America
India
South America
Africa
Indian Ocean
Madagascar
Oceania
Manjekia maturbongsii
Jailoloa halmaherensis
Wallaceodoxa raja ampat
Veitchia winin
Balaka longirostris
Adonidia merrillii
Ponapea hentyi
Ptychococcus lepidotus
Brassiophoenix schumannii
Ptychosperma buabe
Wodyetia bifurcata
Carpentaria acuminata
Normanbya normanbyi
Drymophloeus litigiosus
Chambeyronia macrocarpa
Archontophoenix alexandrae
Actinorhytis calapparia
Calyptrocalyx awa
Cyphophoenix elegans
Burretiokentia vieillardii
Campecarpus fulcitus
Veillonia alba
Physokentia insolita
Lepidorrhachis mooreana
Cyphosperma tanga
Rhopalostylis baueri
Hedyscepe canterburyana
Basselinia moorei
Linospadix palmerianus
Howea forsteriana
Laccospadix australasicus
Heterospathe longipes
Dransfieldia micrantha
Satakentia liukiuensis
Neoveitchia storckii
Carpoxylon macrospermum
Cyphokentia cerifera
Clinosperma vaginata
Hydriastele wendlandiana
Pinanga rupestris
Areca chaiana
Nenga grandiflora
Areca catechu
Bentinckia condapanna
Cyrtostachys renda
Clinostigma ponapense
Tectiphiala ferox
Acanthophoenix rousselii
Deckenia nobilis
Oncosperma fasciculatum
Oncosperma tigillarium
Verschaffeltia splendida
Roscheria melanochaetes
Phoenicophorium borsigianum
Nephrosperma vanhoutteanum
Rhopaloblaste ledermanniana
Dictyosperma album
Iguanura wallichiana
Loxococcus rupicola
Marojejya darianii
Dypsis procumbens
Chrysalidocarpus tanalensis
Vonitra vonitrandambo
Lemurophoenix laevis
Masoala kona
Prestoea ensiformis
Neonicholsonia watsonii
Euterpe edulis
Oenocarpus minor
Hyospathe elegans
Calyptronoma plumeriana
Calyptrogyne tutensis
Geonoma monospatha
Asterogyne martiana
Welfia regia
Pholidostachys dactyloides
Sommieria leucophylla
Pelagodoxa mesocarpa
Manicaria saccifera
Leopoldinia pulchra
Parajubaea torallyi
Allagoptera caudescens
Syagrus cearensis
Jubaea chilensis
Butia capitata
Cocos nucifera
Attalea butyracea
Voanioala gerardii
Jubaeopsis caffra
Beccariophoenix fenestralis
Desmoncus orthacanthos
Bactris panamensis
Astrocaryum alatum
Acrocomia aculeata
Aiphanes eggersii
Elaeis oleifera
Barcella odora
Reinhardtia paiewonskiana
Roystonea regia
Sclerosperma mannii
Orania ravaka
Podococcus barteri
Truongsonia lecongkietii
Gaussia maya
Chamaedorea alternans
Synechanthus warscewiczianus
Wendlandiella gracilis
Hyophorbe lagenicaulis
Socratea exorrhiza
Iriartella setigera
Wettinia aequalis
Iriartea deltoidea
Dictyocaryum lamarckianum
Ravenea albicans
Oraniopsis appendiculata
Juania australis
Ceroxylon quindiuense
Aphandra natalia
Ammandra decasperma
Phytelephas aequatorialis
Pseudophoenix vinifera
Saribus papuanus
Lanonia gracilis
Licuala telifera
Johannesteijsmannia altifrons
Pholidocarpus macrocarpus
Livistona australis
Serenoa repens
Acoelorrhaphe wrightii
Rhapis subtilis
Guihaia argyrata
Maxburretia furtadoana
Trachycarpus fortunei
Chamaerops humilis
Rhapidophyllum hystrix
Brahea edulis
Colpothrinax aphanopetala
Pritchardia arecina
Copernicia prunifera
Washingtonia filifera
Thrinax radiata
Leucothrinax morrisii
Hemithrinax ekmaniana
Zombia antillarum
Coccothrinax argentata
Schippia concolor
Sabinaria magnifica
Itaya amicorum
Chelyocarpus chuco
Trithrinax brasiliensis
Cryosophila bartlettii
Sabal bermudana
Phoenix reclinata
Borassus flabellifer
Borassodendron borneense
Lodoicea maldivica
Latania loddigesii
Medemia argun
Hyphaene coriacea
Satranala decussilvae
Bismarckia nobilis
Corypha umbraculifera
Caryota rumphiana
Arenga undulatifolia
Kerriodoxa elegans
Chuniophoenix nana
Tahina spectabilis
Nannorrhops ritchieana
Nypa fruticans
Plectocomia mulleri
Myrialepis paradoxa
Plectocomiopsis geminiflora
Calamus calospathus
Pigafetta filaris
Metroxylon sagu
Salacca bakeriana
Eleiodoxa conferta
Korthalsia echinometra
Eugeissona tristis
Oncocalamus tuleyi
Laccosperma opacum
Eremospatha cuspidata
Raphia farinifera
Mauritiella armata
Mauritia flexuosa
Lepidocaryum tenue
Ma
100
66
56
34
23
0
Early Cretaceous
Late Cretaceous
Paleocene
Eocene
Oligocene
Neogene +
Quaternary

Figure S7

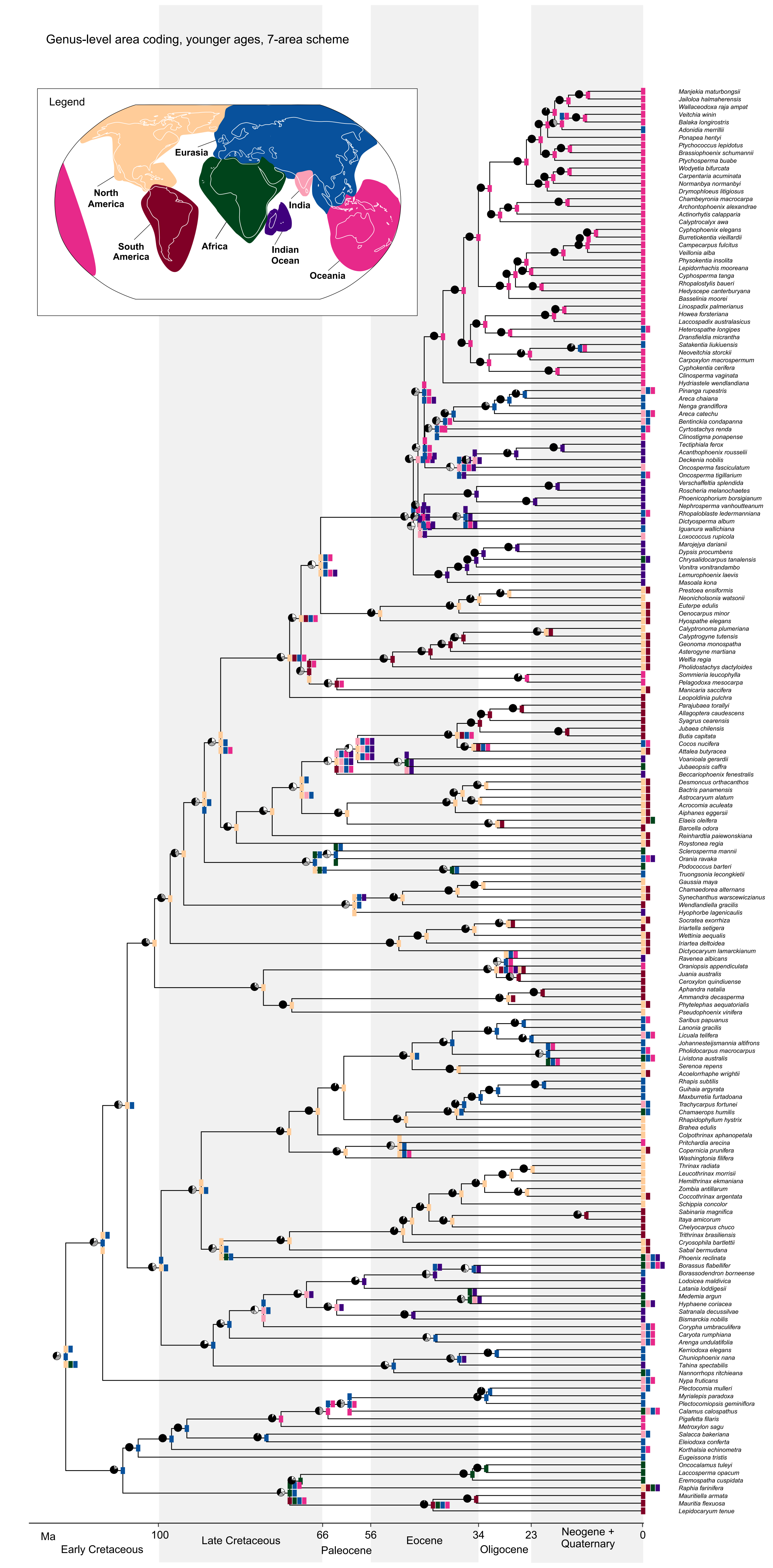
Genus-level area coding, younger ages, 7-area scheme
Legend
Eurasia
North America
South America
Africa
India
Indian Ocean
Oceania
Manjekia maturbongsii
Jailoloa halmaherensis
Wallaceodoxa raja ampat
Veitchia winin
Balaka longirostris
Adonidia merrillii
Ponapea hentyi
Ptychococcus lepidotus
Brassiophoenix schumannii
Ptychosperma buabe
Wodyetia bifurcata
Carpentaria acuminata
Normanbya normanbyi
Drymophloeus litigiosus
Chambeyronia macrocarpa
Archontophoenix alexandrae
Actinorhytis calapparia
Calyptrocalyx awa
Cyphophoenix elegans
Burretiokentia vieillardii
Campecarpus fulcitus
Veillonia alba
Physokentia insolita
Lepidorrhachis mooreana
Cyphosperma tanga
Rhopalostylis baueri
Hedyscepe canterburyana
Basselinia moorei
Linospadix palmerianus
Howea forsteriana
Laccospadix australasicus
Heterospathe longipes
Dransfieldia micrantha
Satakentia liukiuensis
Neoveitchia storckii
Carpoxylon macrospermum
Cyphokentia cerifera
Clinosperma vaginata
Hydriastele wendlandiana
Pinanga rupestris
Areca chaiana
Nenga grandiflora
Areca catechu
Bentinckia condapanna
Cyrtostachys renda
Clinostigma ponapense
Tectiphiala ferox
Acanthophoenix rousselii
Deckenia nobilis
Oncosperma fasciculatum
Oncosperma tigillarium
Verschaffeltia splendida
Roscheria melanochaetes
Phoenicophorium borsigianum
Nephrosperma vanhoutteanum
Rhopaloblaste ledermanniana
Dictyosperma album
Iguanura wallichiana
Loxococcus rupicola
Marojejya darianii
Dypsis procumbens
Chrysalidocarpus tanalensis
Vonitra vonitrandambo
Lemurophoenix laevis
Masoala kona
Prestoea ensiformis
Neonicholsonia watsonii
Euterpe edulis
Oenocarpus minor
Hyospathe elegans
Calyptronoma plumeriana
Calyptrogyne tutensis
Geonoma monospatha
Asterogyne martiana
Welfia regia
Pholidostachys dactyloides
Sommieria leucophylla
Pelagodoxa mesocarpa
Manicaria saccifera
Leopoldinia pulchra
Parajubaea torallyi
Allagoptera caudescens
Syagrus cearensis
Jubaea chilensis
Butia capitata
Cocos nucifera
Attalea butyracea
Voanioala gerardii
Jubaeopsis caffra
Beccariophoenix fenestralis
Desmoncus orthacanthos
Bactris panamensis
Astrocaryum alatum
Acrocomia aculeata
Aiphanes eggersii
Elaeis oleifera
Barcella odora
Reinhardtia paiewonskiana
Roystonea regia
Sclerosperma mannii
Orania ravaka
Podococcus barteri
Truongsonia lecongkietii
Gaussia maya
Chamaedorea alternans
Synechanthus warscewiczianus
Wendlandiella gracilis
Hyophorbe lagenicaulis
Socratea exorrhiza
Iriartella setigera
Wettinia aequalis
Iriartea deltoidea
Dictyocaryum lamarckianum
Ravenea albicans
Oraniopsis appendiculata
Juania australis
Ceroxylon quindiuense
Aphandra natalia
Ammandra decasperma
Phytelephas aequatorialis
Pseudophoenix vinifera
Saribus papuanus
Lanonia gracilis
Licuala telifera
Johannesteijsmannia altifrons
Pholidocarpus macrocarpus
Livistona australis
Serenoa repens
Acoelorrhaphe wrightii
Rhapis subtilis
Guihaia argyrata
Maxburretia furtadoana
Trachycarpus fortunei
Chamaerops humilis
Rhapidophyllum hystrix
Brahea edulis
Colpothrinax aphanopetala
Pritchardia arecina
Copernicia prunifera
Washingtonia filifera
Thrinax radiata
Leucothrinax morrisii
Hemithrinax ekmaniana
Zombia antillarum
Coccothrinax argentata
Schippia concolor
Sabinaria magnifica
Itaya amicorum
Chelyocarpus chuco
Trithrinax brasiliensis
Cryosophila bartlettii
Sabal bermudana
Phoenix reclinata
Borassus flabellifer
Borassodendron borneense
Lodoicea maldivica
Latania loddigesii
Medemia argun
Hyphaene coriacea
Satranala decussilvae
Bismarckia nobilis
Corypha umbraculifera
Caryota rumphiana
Arenga undulatifolia
Kerriodoxa elegans
Chuniophoenix nana
Tahina spectabilis
Nannorrhops ritchieana
Nypa fruticans
Plectocomia mulleri
Myrialepis paradoxa
Plectocomiopsis geminiflora
Calamus calospathus
Pigafetta filaris
Metroxylon sagu
Salacca bakeriana
Eleiodoxa conferta
Korthalsia echinometra
Eugeissona tristis
Oncocalamus tuleyi
Laccosperma opacum
Eremospatha cuspidata
Raphia farinifera
Mauritiella armata
Mauritia flexuosa
Lepidocaryum tenue
Ma
100
66
56
34
23
0
Early Cretaceous
Late Cretaceous
Paleocene
Eocene
Oligocene
Neogene + Quaternary

Figure S8

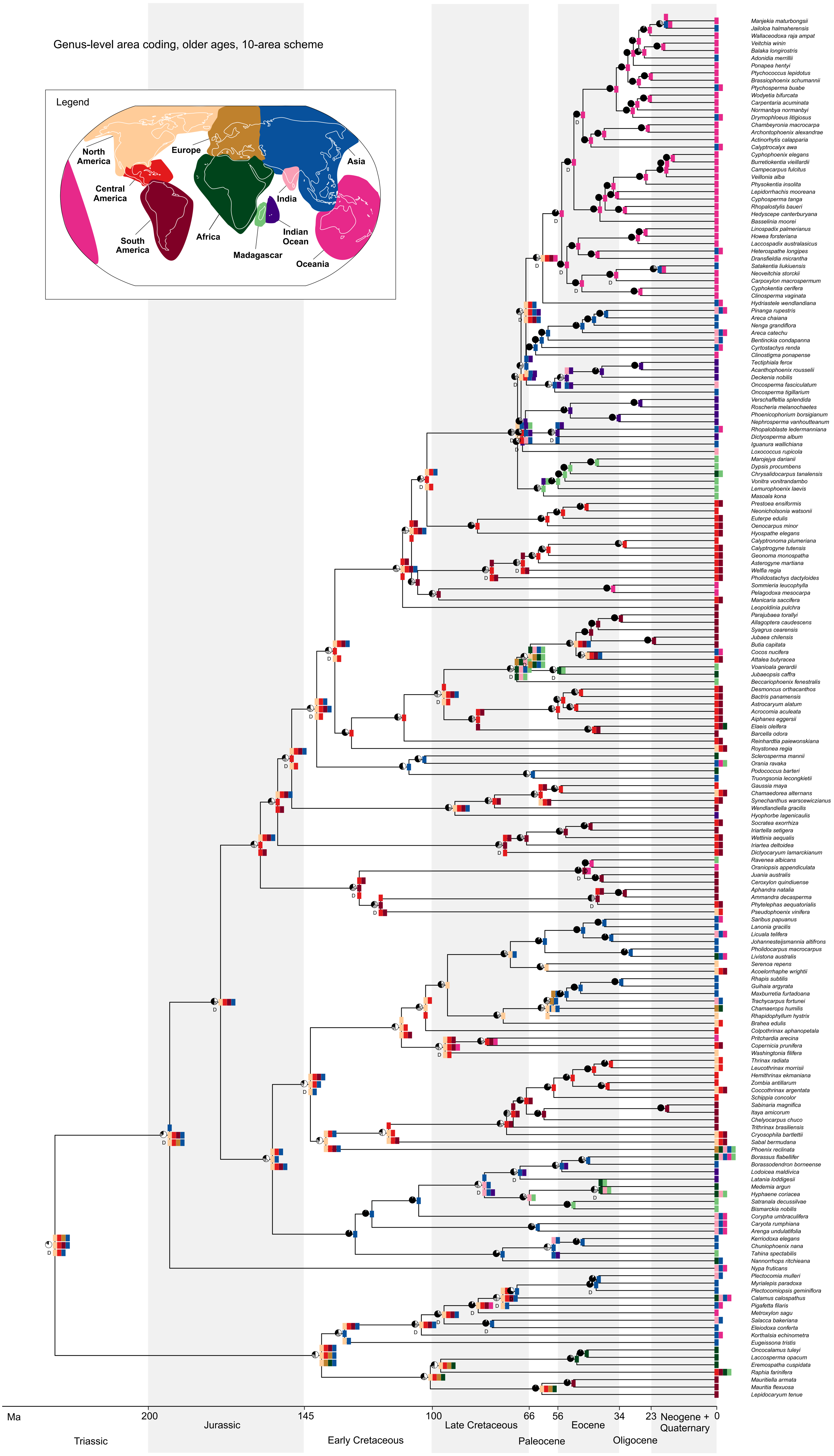

Genus-level area coding, older ages, 10-area scheme
Legend
North America
Central America
South America
Europe
Africa
Madagascar
India
Indian Ocean
Asia
Oceania
Manjekia maturbongsii
Jailoloa halmaherensis
Wallaceodoxa raja ampat
Veitchia winin
Balaka longirostris
Adonidia merrillii
Ponapea hentyi
Ptychococcus lepidotus
Brassiophoenix schumannii
Ptychosperma buabe
Wodyetia bifurcata
Carpentaria acuminata
Normanbya normanbyi
Drymophloeus litigiosus
Chambeyronia macrocarpa
Archontophoenix alexandrae
Actinorhytis calapparia
Calyptrocalyx awa
Cyphophoenix elegans
Burretiokentia vieillardii
Campecarpus fulcitus
Veillonia alba
Physokentia insolita
Lepidorrhachis mooreana
Cyphosperma tanga
Rhopalostylis baueri
Hedyscepe canterburyana
Basselinia moorei
Linospadix palmerianus
Howea forsteriana
Laccospadix australasicus
Heterospathe longipes
Dransfieldia micrantha
Satakentia liukiuensis
Neoveitchia storckii
Carpoxylon macrospermum
Cyphokentia cerifera
Clinosperma vaginata
Hydriastele wendlandiana
Pinanga rupestris
Areca chaiana
Nenga grandiflora
Areca catechu
Bentinckia condapanna
Cyrtostachys renda
Clinostigma ponapense
Tectiphiala ferox
Acanthophoenix rousselii
Deckenia nobilis
Oncosperma fasciculatum
Oncosperma tigillarium
Verschaffeltia splendida
Roscheria melanochaetes
Phoenicophorium borsigianum
Nephrosperma vanhoutteanum
Rhopaloblaste ledermanniana
Dictyosperma album
Iguanura wallichiana
Loxococcus rupicola
Marojejya darianii
Dypsis procumbens
Chrysalidocarpus tanalensis
Vonitra vonitrandambo
Lemurophoenix laevis
Masoala kona
Prestoea ensiformis
Neonicholsonia watsonii
Euterpe edulis
Oenocarpus minor
Hyospathe elegans
Calyptronoma plumeriana
Calyptrogyne tutensis
Geonoma monospatha
Asterogyne martiana
Welfia regia
Pholidostachys dactyloides
Sommieria leucophylla
Pelagodoxa mesocarpa
Manicaria saccifera
Leopoldinia pulchra
Parajubaea torallyi
Allagoptera caudescens
Syagrus cearensis
Jubaea chilensis
Butia capitata
Cocos nucifera
Attalea butyracea
Voanioala gerardii
Jubaeopsis caffra
Beccariophoenix fenestralis
Desmoncus orthacanthos
Bactris panamensis
Astrocaryum alatum
Acrocomia aculeata
Aiphanes eggersii
Elaeis oleifera
Barcella odora
Reinhardtia paiewonskiana
Roystonea regia
Sclerosperma mannii
Orania ravaka
Podococcus barteri
Truongsonia lecongkietii
Gaussia maya
Chamaedorea alternans
Synechanthus warscewiczianus
Wendlandiella gracilis
Hyophorbe lagenicaulis
Socratea exorrhiza
Iriartella setigera
Wettinia aequalis
Iriartea deltoidea
Dictyocaryum lamarckianum
Ravenea albicans
Oraniopsis appendiculata
Juania australis
Ceroxylon quindiuense
Aphandra natalia
Ammandra decasperma
Phytelephas aequatorialis
Pseudophoenix vinifera
Saribus papuanus
Lanonia gracilis
Licuala telifera
Johannesteijsmannia altifrons
Pholidocarpus macrocarpus
Livistona australis
Serenoa repens
Acoelorrhaphe wrightii
Rhapis subtilis
Guihaia argyrata
Maxburretia furtadoana
Trachycarpus fortunei
Chamaerops humilis
Rhapidophyllum hystrix
Brahea edulis
Colpothrinax aphanopetala
Pritchardia arecina
Copernicia prunifera
Washingtonia filifera
Thrinax radiata
Leucothrinax morrisii
Hemithrinax ekmaniana
Zombia antillarum
Coccothrinax argentata
Schippia concolor
Sabinaria magnifica
Itaya amicorum
Chelyocarpus chuco
Trithrinax brasiliensis
Cryosophila bartlettii
Sabal bermudana
Phoenix reclinata
Borassus flabellifer
Borassodendron borneense
Lodoicea maldivica
Latania loddigesii
Medemia argun
Hyphaene coriacea
Satranala decussilvae
Bismarckia nobilis
Corypha umbraculifera
Caryota rumphiana
Arenga undulatifolia
Kerriodoxa elegans
Chuniophoenix nana
Tahina spectabilis
Nannorrhops ritchieana
Nypa fruticans
Plectocomia mulleri
Myrialepis paradoxa
Plectocomiopsis geminiflora
Calamus calospathus
Pigafetta filaris
Metroxylon sagu
Salacca bakeriana
Eleiodoxa conferta
Korthalsia echinometra
Eugeissona tristis
Oncocalamus tuleyi
Laccosperma opacum
Eremospatha cuspidata
Raphia farinifera
Mauritiella armata
Mauritia flexuosa
Lepidocaryum tenue
Ma
200
145
100
66
56
34
23
0
Triassic
Jurassic
Early Cretaceous
Late Cretaceous
Paleocene
Eocene
Oligocene
Neogene + Quaternary
